# *Phf15*—a novel transcriptional repressor regulating inflammation in mouse microglia

**DOI:** 10.1101/2019.12.17.879940

**Authors:** Sandra E. Muroy, Greg A. Timblin, Marcela K. Preininger, Paulina Cedillo, Kaoru Saijo

## Abstract

**Aim:** Excessive microglial inflammation has emerged as a key player in mediating the effects of aging and neurodegeneration on brain dysfunction. Thus, there is great interest in discovering transcriptional repressors that can control this process. We aimed to examine whether *Phf15*—one of the top differentially expressed genes in microglia during aging in humans—could regulate transcription of pro-inflammatory mediators in microglia.

**Methods:** RT-qPCR was used to assess *Phf15* mRNA expression in mouse brain during aging. Loss-of-function (shRNA-mediated knockdown (KD) and CRISPR/Cas9-mediated knockout (KO) of *Phf15)* and gain-of-function (retroviral overexpression (OE) of murine *Phf15* cDNA) studies in a murine microglial cell line (SIM-A9) followed by immune activation with lipopolysaccharide (LPS) were used to determine the effect of *Phf15* on pro-inflammatory factor (*Tnfα, Il-1β, Nos2*) mRNA expression. RNA-sequencing was used to determine global transcriptional changes after *Phf15* knockout under basal conditions and after LPS stimulation.

**Results:** *Phf15* expression increases in mouse brain during aging, similar to humans. KD, KO and OE studies determined that *Phf15* represses mRNA expression levels of pro-inflammatory mediators such as *Tnfα, Il-1β* and *Nos2*. Global transcriptional changes after *Phf15* KO showed that *Phf15* specifically represses genes related to the antiviral (type I interferon) response and cytokine production in microglia.

**Conclusion:** We provide the first evidence that *Phf15* is an important transcriptional repressor of microglial inflammation, regulating the antiviral response and pro-inflammatory cytokine production. Importantly, *Phf15* regulates both basal and signal-dependent activation and controls the magnitude and duration of the microglial inflammatory response.

## Introduction

Microglia are the resident myeloid-lineage cells of the brain. They actively provide homeostatic surveillance of the brain parenchyma playing critical roles during development, maintenance and repair throughout the life of an organism. As innate immune cells, however, microglia are also capable of mounting a full inflammatory response to environmental challenge in order to clear threats and restore homeostasis^*[1–7]*^. Microglia express pattern recognition receptors including Toll-like receptors (TLRs) to sense changes in their environment, such as infection by pathogens or endogenous danger signals. They can then respond by releasing pro-inflammatory mediators such as Tumor necrosis factor alpha (TNFα), Interleukin 1 beta (IL-1β), Interleukin 6 (IL-6), reactive oxygen species (ROS) and reactive nitrogen species (RNS) including nitric oxide (NO) to protect against threats^*[3,5,7]*^.

Although beneficial when their production is tightly controlled, deregulated or sustained microglial production of inflammatory mediators can lead to collateral damage of surrounding neurons and other cells^*[5,7,8]*^. Thus, the transition to an activated state, as well as, timely resolution of the inflammatory response, must be tightly regulated. Increasing evidence suggests that during aging, microglia lose homeostatic function and acquire a pro-inflammatory phenotype that exacerbates aging-related brain dysfunction^*[9]*^. Indeed, aberrant microglia activation has been found in many types of age-related neurodegenerative conditions for example, Parkinson’s disease (PD) and Alzheimer’s disease (AD) which are marked by inflammatory processes involving glia, and microglia in particular^*[9–11]*^.

Since excessive production of pro-inflammatory mediators is neurotoxic^*[8,12–14]*^, various molecular mechanisms exist to regulate transcriptional repression of inflammatory gene expression. For example, basal state repression, that is, before the arrival of an activating signal, is generally carried out via recruitment of co-repressor complexes that prevent initiation of inflammatory gene transcription. After stimulation by an activating signal, additional mechanisms can maintain quiescence by restraining active transcription. Finally, numerous mechanisms mediate the timely resolution of the inflammatory response at the transcriptional level, including transrepression mechanisms that can remove transcription factors from inflammatory gene promoters^*[8,15–18]*^.

Studies have also highlighted an important role for chromatin modifications in the transcriptional control of inflammatory gene expression^*[19,20]*^. A recent study by Soreq et al.^*[21]*^, which compared transcriptional profiles of different brain cell types and regions throughout healthy human aging found microglial gene expression profiles as being one of the most predictive markers of biological age in the brain^*[21]*^. The same study identified a relatively unknown gene, PHD finger protein 15 *(PHF15)* among the top 25 differentially expressed genes in microglia during aging. Work in embryonic stem cells, and sequence and structural similarity to other members of the PHF family, indicate that *PHF15* is a putative chromatin-mediated gene regulator^*[22,23]*^.

Given that aging skews microglia towards a pro-inflammatory phenotype, and that *PHF15* was found to be highly upregulated during non-pathological aging, we sought to determine whether *Phf15* might regulate microglial inflammatory function. We found that *Phf15* strongly represses pro-inflammatory gene expression, regulating both basal and signal-dependent activation and modulating the magnitude and duration of the mouse microglial inflammatory response. Importantly, *Phf15* seems to regulate pro-inflammatory and Interferon type I (IFN-I)-dependent gene expression. Increased IFN-I tone and pro-inflammatory cytokine expression are both hallmarks of the aging brain^*[24–28]*^. Our findings suggest that *Phf15* is an important novel repressor of microglial inflammatory function that might work to counteract age-induced inflammation in the healthy, aging brain.

## METHODS

### Animals

Adult male C57Bl6/J mice were purchased from The Jackson Laboratory and maintained on a 12:12-h light–dark cycle (lights on at 0700 hours) with *ad libitum* access to food and water and aged for ~2.5, ~14 or ~20 months. All animal care and procedures were approved by the University of California, Berkeley Animal Care and Use Committee.

### shRNA-mediated knockdown of *Phf15* in murine microglial cells

pGIPZ Lentiviral mouse Jade2 shRNA constructs or a control scrambled shRNA were purchased from Dharmacon (Lafayette, CO). Lentivirus was packaged via co-transfection of each pGIPZ-shRNA with pCMV-VSV-G (Addgene plasmid #8454)^[29]^ and pCMV-dR8.2 (Addgene plasmid #8455)^[29]^ into HEK 293T cells using Lipofectamine 3000 reagent (Life Technologies, Carlsbad, CA) according to the manufacturer’s instructions. Viral supernatant was harvested after 48 hours and incubated with SIM-A9 murine microglial cells in SIM-A9 complete medium (DMEM/F12, Life Technologies, Carlsbad, CA), 10% fetal bovine serum (FBS; GE Healthcare Life Sciences, Chicago, IL), 5% horse serum (HS; GE Healthcare Life Sciences, Chicago, IL), 1% Pen-Strep (Life Technologies, Carlsbad, CA)). After 48 hours, GFP+ cells were sorted by FACS on an Aria Fusion (BD Biosciences, San Jose, CA; UC Berkeley Cancer Research Laboratory), expanded and subcultured for immune stimulation experiments. Percent knockdown was determined via RT-qPCR.

### Overexpression of *Phf15* in murine microglial cells

A *Phf15* overexpression vector was constructed by cloning the full length *Phf15* cDNA (Mus *musculus* PHD finger protein 15, mRNA cDNA clone MGC: 143877 IMAGE:40094330) obtained from Dharmacon (Lafayette. CO) into a pMYs-IRES-GFP retroviral vector (Cell Biolabs Inc, San Diego, CA). Virus expressing the full length *Phf15* cDNA or empty vector control were co-transfected with pCL-10 A1 (Addgene plasmid #15805)^[30]^ in HEK293T cells using Lipofectamine 3000 (Life Technologies, Carlsbad, CA) reagent according to the manufacturer’s instructions. SIM-A9 cells were incubated with virus for 24 hrs and then sorted via FACS on an Aria Fusion, expanded and subcultured for immune stimulation experiments. Fold overexpression was verified via RT-qPCR.

### Generation of *Phf15* knockout microglia

*Phf15* knockout (KO) SIM-A9 cells were generated using the Alt-R CRISPR–Cas9–mediated gene editing system (guide RNA sequence ACTACATCCTGGCGGACCCGTGG) from IDT (Coralville, IA) using CRISPRMAX Lipofectamine reagent (IDT) as per the manufacturer’s instructions. ATTO 550+ cells were single-cell sorted on an Aria Fusion. Clones were screened for *Phf15* deletion using PCR (primers Forward: agcacacttgtaaccctcct and Reverse: gaccaatgtctgttgttgttcg) followed by restriction digest with BtgI (New England Biolabs, Ipswich, MA). Percent decrease in *Phf15* mRNA transcript expression was determined via RT-qPCR (primer sequences are listed in Supplementary Table 1).

### Immune stimulation

For all immune stimulation time course experiments, cells (knockdown, knockout, overexpression and respective controls) were subcultured in 24-well plates at a density of 10^5^ cells/well (in triplicate) and stimulated with LPS (final concentration of 100 ng/ml; Sigma Aldrich, St. Louis, MO), CpG ODN (final concentration of 2.5 uM; InvivoGen, San Diego, CA) or Poly(I:C) (final concentration of 25 uM; Sigma Aldrich, St. Louis, MO) for 1, 6, 12 or 24 hrs. No stimulation controls received an equivalent volume of sterile 1xPBS (Invitrogen, Carlsbad, CA).

### RNA extraction

Mice were sacrificed according to the approved protocol. Brains were isolated and frontal cortical areas were dissected, flash frozen and stored at - 80 °C. RNA was extracted using a bead homogenizer (30 seconds, setting ‘5’’; Bead Mill, VWR) in Trizol reagent (ThermoFisher, Waltham, MA). Total RNA was extracted using the Direct-zol RNA miniprep kit (Zymo Research, Irvine, CA) according to the manufacturer’s instructions. For cell lines, after immune stimulation, media was aspirated and wells were washed 2x with ice-cold 1xPBS (Invitrogen, Carlsbad, CA). RNA was extracted using the Direct-zol RNA miniprep kit (Zymo Research, Irvine, CA).

### Real-Time Quantitative PCR (RT-qPCR)

cDNA was reversed transcribed from total RNA using the SuperScript™ III First-Strand Synthesis System kit (ThermoFisher, Waltham, MA) following the manufacturer’s instructions. RT-qPCR was run using SYBR green (Roche) on a QuantStudio 6 (ThermoFisher, Waltham, MA) real-time PCR machine. All RT-qCPR primers are specific to the desired template, span exon-exon junctions and capture all transcript variants for the specific gene under study. Ct values were normalized to the housekeeping gene *Hprt*. Primer sequences used in this study are listed in Supplementary Table 1.

### RNA-seq library preparation and analysis

RNA was extracted from a total of n= 3 replicates per condition (*Phf15* KO or control) and was used to prepare libraries for RNA sequencing using the KAPA mRNA HyperPrep Kit according to the manufacturer’s instructions (KAPA Biosystems, Wilmington, MA). Libraries were quality control checked via Qubit (ThermoFisher, Waltham, MA) and via RT-qPCR with a next generation sequencing (NGS) library quantification kit (Zymo Research, Irvine, CA). RNA sequencing (1 lane) was performed on a HiSeq4000 sequencing system (Illumina Inc., San Diego, CA; UC Berkeley Genomics Sequencing Laboratory). Sequencing reads were aligned to the *Mus musculus* reference genome assembly GRCm38 (mm10) using Spliced Transcripts Alignment to a Reference (STAR) aligner^[31]^. Count data was analyzed with Hypergeometric Optimization of Motif EnRichment (HOMER) software for next-generation sequencing analysis (http://homer.ucsd.edu/homer/ngs/index.html) which uses the R/Bioconductor package DESeq2^[32]^ to perform differential gene expression analysis. To adjust for multiple comparisons, DESeq2 uses the Benjamini-Hochberg procedure to control the false discovery rate (FDR) and returned FDR adjusted *p* values and log2fold expression changes between *Phf15* KO and control conditions for each gene. Genes were filtered by adjusted *p* value (adjusted p < 0.01 for upregulated genes or 0.05 for downregulated genes) and log2 fold change in expression (greater than 1.5 log2fold change for upregulated genes and less than −1.5 for downregulated genes). Too few downregulated genes (< 200) passed the more stringent adjusted p < 0.01 cutoff for robust downstream biological function analysis, so the adjusted *p* value threshold was lowered to *p* < 0.05. Results were visualized using the R package EnhancedVolcano^[33]^. Lists of upregulated and downregulated genes were input into Metascape^[34]^, a gene annotation and analysis tool, to determine enriched biological themes within the gene lists.

### Motif enrichment

Transcription factor binding sites (‘motifs’) were analyzed using HOMER (http://homer.ucsd.edu/homer/ngs/index.html).

### Statistical Analysis

Relative mRNA expression of *Phf15* in mouse frontal cortical areas was analyzed using ordinary one-way ANOVA with post hoc Tukey’s multiple comparisons to compare expression levels across age. Percent knockdown and time course experiments measuring expression levels of inflammatory markers (*Tnfα, Nos2, Il-1β*) between control and *Phf15* shRNAs shPhf15-1 and shPhf15-2 after immune stimulation (with LPS, CpG-ODN or Poly(I:C)) were analyzed via Unpaired t-tests between each shRNA versus control shRNA within timepoint. Fold overexpression or percent reduction and time course experiments for *Phf15* overexpression and knockout cell lines were analyzed using Unpaired t-tests (overexpression or knockout vs. respective control) within each time point. *P* < 0.05 was considered significant in all experiments.

## RESULTS

### Aging increases *Phf15* expression in mouse brain

To investigate whether *Phf15* increases in mouse brains similar to humans^[21]^, we measured *Phf15* mRNA expression in mouse frontal cortical brain areas across age. We were interested in frontal cortical regions because of their involvement in mediating various aspects of cognitive function and because they are selectively affected in several aging-related neurodegenerative conditions like PD, AD and frontotemporal dementia (FTD)^[35,36]^.

We found that compared to young (~2.5-month-old) mice, old (~20-month-old) mice had significantly elevated *Phf15* mRNA levels in frontal cortical areas (Figure 1). Middle-aged (~14-month-old) mice showed a trend towards increased *Phf15* mRNA expression that did not reach statistical significance. Our data suggest that *Phf15* expression increases in mouse frontal cortical regions upon normal aging, similar to what was previously reported in humans^*[21]*^.

**Figure 1.**
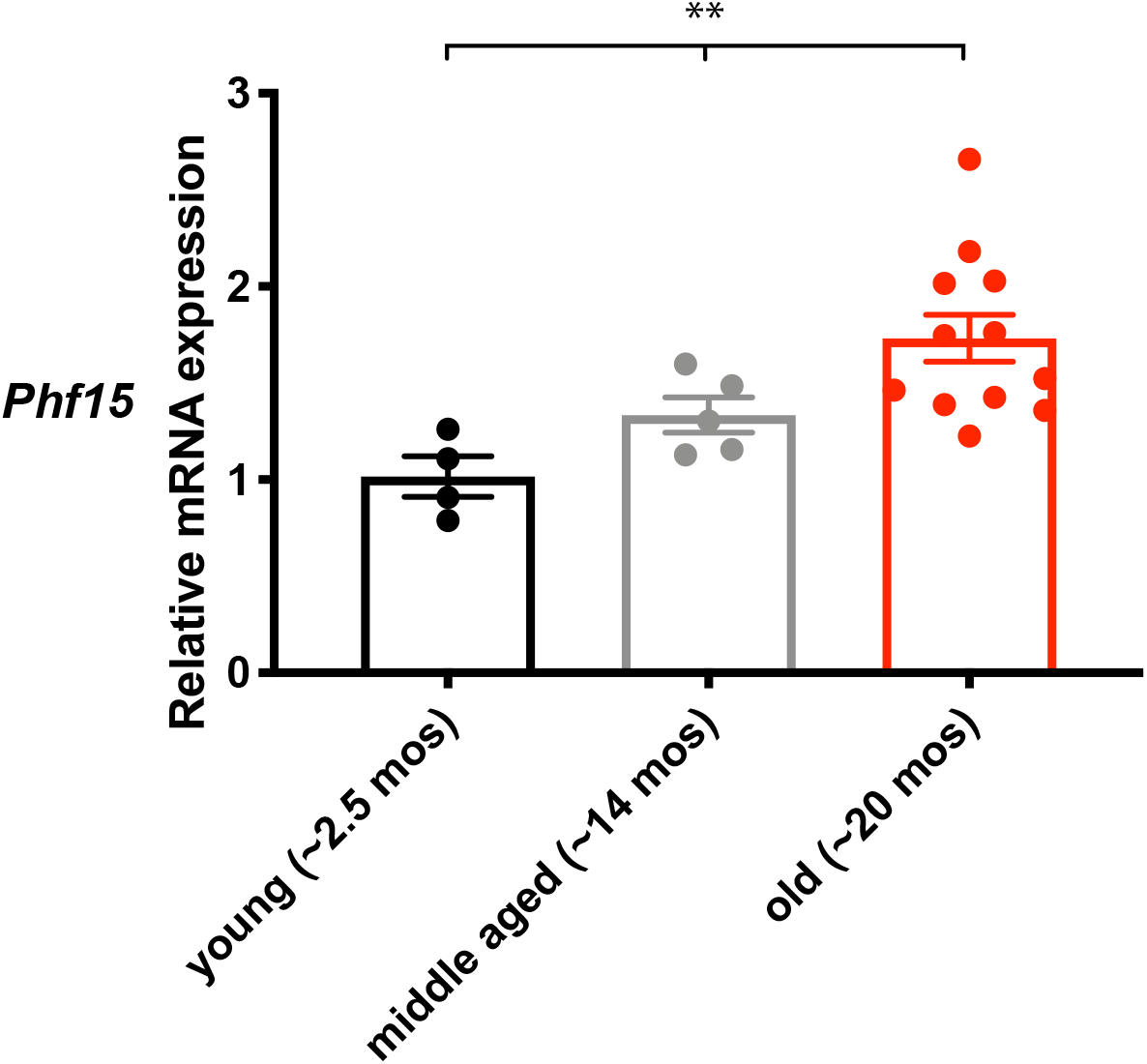
*Phf15* expression increases in aged mouse frontal cortical areas. *Phf15* mRNA expression was significantly elevated in frontal cortical areas of old (~20-month-old; red bar) mice compared to young (~2.5 month old; black bar) mice. Data are mean ±SEM (n = 4 young, n = 5 middle aged, n = 12 old). One-way ANOVA with Tukey’s post hoc comparisons between age groups: \*\**p*<0.01

### Knockdown of *Phf15* increases the magnitude of the microglial inflammatory response

To determine whether *Phf15* regulates microglial inflammatory function, we performed loss-of function studies via shRNA-mediated knockdown (KD) in a murine microglial cell line, SIM-A9, followed by immune activation with lipopolysaccharide (LPS), a component of gram-negative bacterial cell walls and Toll-like receptor 4 (TLR4) agonist. We chose LPS because 1) Intraperitoneal and/or intracranial administration of LPS in mice lead to increased microglial activation, neuroinflammation, neuronal loss including loss of dopaminergic neurons in the substantia nigra in a mouse model of PD^*[8]*^ as well as, cognitive and neurological deficits^[37]^, 2) Aged individuals show increased systemic levels of LPS in the bloodstream^[38]^ which are associated with increased inflammation and microglial activation^[39]^ and 3) In humans, TLR4 activation is linked to age-related pathologies like PD and AD^[40–42]^. Thus, LPS serves as a relevant aging-related physiological immune stimulant.

KD of *Phf15* resulted in a significant reduction in *Phf15* mRNA transcript levels of 52% or 60% for cell lines shPhf15-1 or shPhf15-2, respectively (Figure 2A), as well as, significantly increased mRNA expression of *Tnfα*, a pro-inflammatory cytokine, after KD with shPhf15-2 at 0, 1, 6 and 12 hours after LPS stimulation (Figure 2B). Similarly, mRNA levels of *Nos2*, the enzyme that catalyzes the production of NO, were significantly elevated at 1, 6 and 12 hours post stimulation for shPhf15-2 and 0, 6 and 12 hours for shPhf15-1 (Figure 2D). Overall, our experiments show that ~50-60% KD, the equivalent of a “heterozygous” condition, results in increased expression of pro-inflammatory mediators over a 12 hour time course that resolves and falls below control levels by 24 hours after immune stimulation. Importantly, microglial inflammatory function was elevated in the absence of immune stimulation (0 hour time point, Figures 2B, D and No stimulation condition, Figures 2C, E), suggesting a loss of repressive mechanisms that inhibit basal state inflammatory gene transcription.

**Figure 2.**
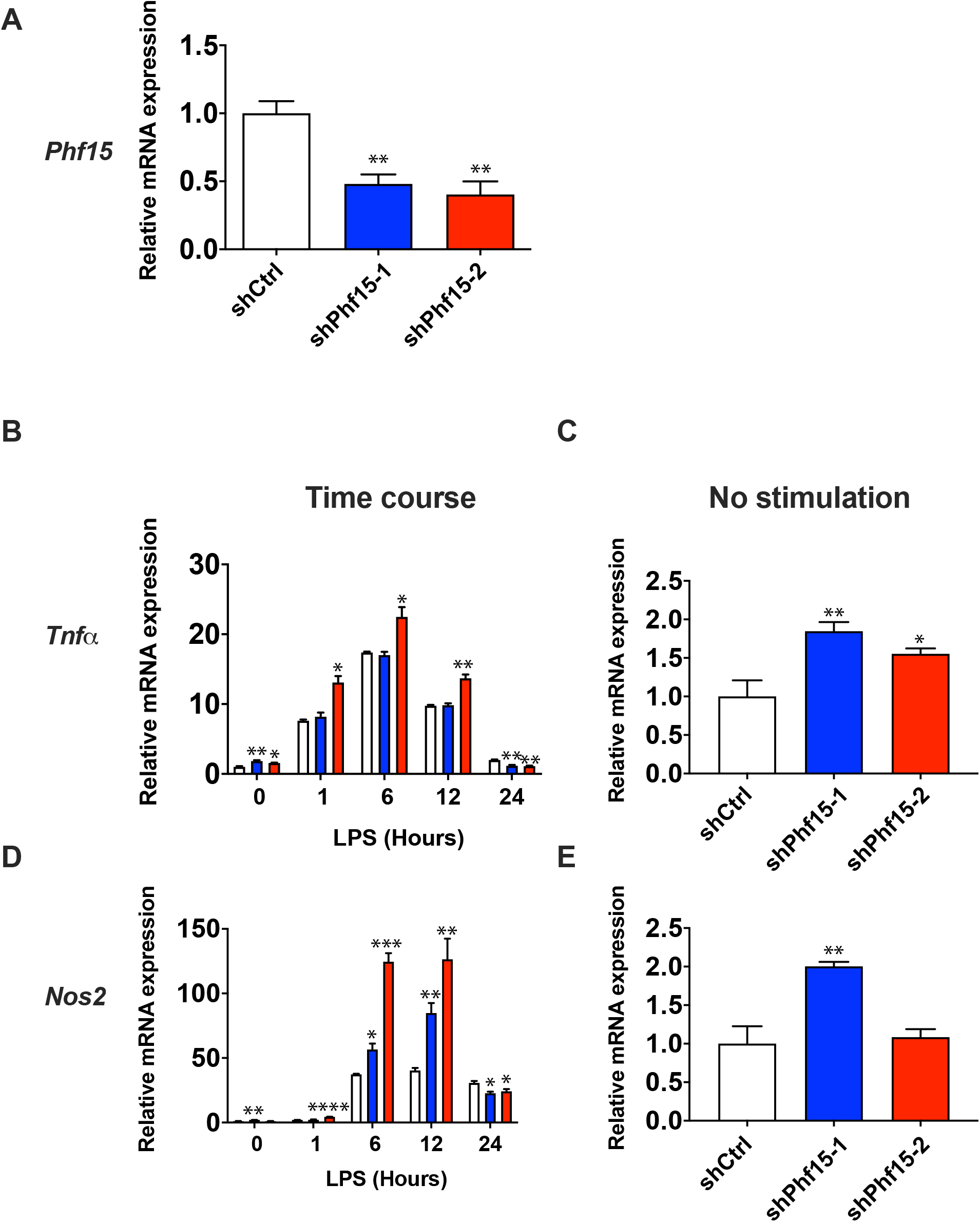
Knockdown of *Phf15* increases the magnitude of the microglial inflammatory response. (A) Knockdown efficiency for anti-Phf15 shRNAs *shPhf15-1* (blue bar, 52% knockdown) and sh*Phf15-2* (red bar, 60% knockdown) Data are mean ±SEM (n = 3 per condition). Unpaired t-tests between sh*Phf15-1* or sh*Phf15-2* and shCtrl cells: asterisks indicate \*\**p*<0.01. 24-hour time course experiments showing relative mRNA expression levels of *Tnfα* (B) and *Nos2* (D) after LPS stimulation of shRNAs sh*Phf15-1* and sh*Phf15-2* compared to shCtrl (control scrambled shRNA). “No stimulation” 0 hr time point is shown for *Tnfα* (C) and *Nos2* (E). Data are mean ±SEM (n = 3 per condition). Unpaired t-tests for sh*Phf15-1* or sh*Phf15-2* and shCtrl cells for individual timepoints: asterisks indicate \**p*<0.05, \*\**p*<0.01, \*\*\**p*<0.001, \*\*\*\**p*<0.0001. LPS, lipopolysaccharide; *Tnfα*, tumor necrosis factor alpha; *Nos2*, nitric oxide synthase, inducible.

We repeated the immune activation time course experiments in *Phf15* KD cells using two separate immune stimulants specific to two distinct TLRs to test the pathway specificity of the inflammatory response. CpG Oligodeoxynucleotide (CpG ODN), a synthetic bacterial and viral DNA mimic targets TLR9 ligand and Polyinosinic:polycytidylic acid (Poly(I:C)), a synthetic viral dsRNA mimic targets TLR3. Although TLR4 uses both the Myeloid differentiation primary response 88 (MyD88) and TIR-domain-containing adapter-inducing interferon-β (TRIF) downstream adapters to transduce its inflammatory cascade, TLR9 and TLR3 utilize MyD88 *or* TRIF respectively (Supplementary Figure 1)^[43,44]^.

Immune stimulation with CpG ODN and Poly(I:C) both yielded similar results to those obtained with LPS stimulation (Supplementary Figures 2 and 3, respectively) denoting no adapter selectivity and confirming that *Phf15* antagonizes gene expression downstream of *both* the MyD88 and TRIF signaling pathways.

### Genetic deletion of *Phf15* increases the magnitude *and* prolongs the duration of the microglial inflammatory response

Since our KD strategy resulted in ~50% reduction in *Phf15* mRNA expression, we next performed CRISPR/Cas9-mediated genetic deletion of *Phf15* in SIM-A9 microglial cells followed by immune activation with LPS. Knockout (KO) of *Phf15* (Figure 3A) resulted in significantly increased LPS-induced expression of *Tnfα* (Figure 3B), *Il-1β* (Figure 3D) and *Nos2*, albeit to a lesser extent (Figure 3F) over a 24-hour time course. Importantly, mRNA levels of both *Tnfα* and *Il-1β* remained elevated at 24 hours compared to control cells, denoting a prolonged inflammatory response and failure to return to steady-state. mRNA expression of *Nos2* showed a significant upregulation over 12 hours (0, 1 and 12 hour timepoints) but had returned to control levels by 24 hours (Figure 3F). Notably, basal expression of all 3 genes was significantly elevated, with a 4-fold increase in *Tnfα*, 14-fold increase in *Il-1β* and 32-fold increase in *Nos2* when comparing KO versus control cells Figures 3C, E, G).

**Figure 3.**
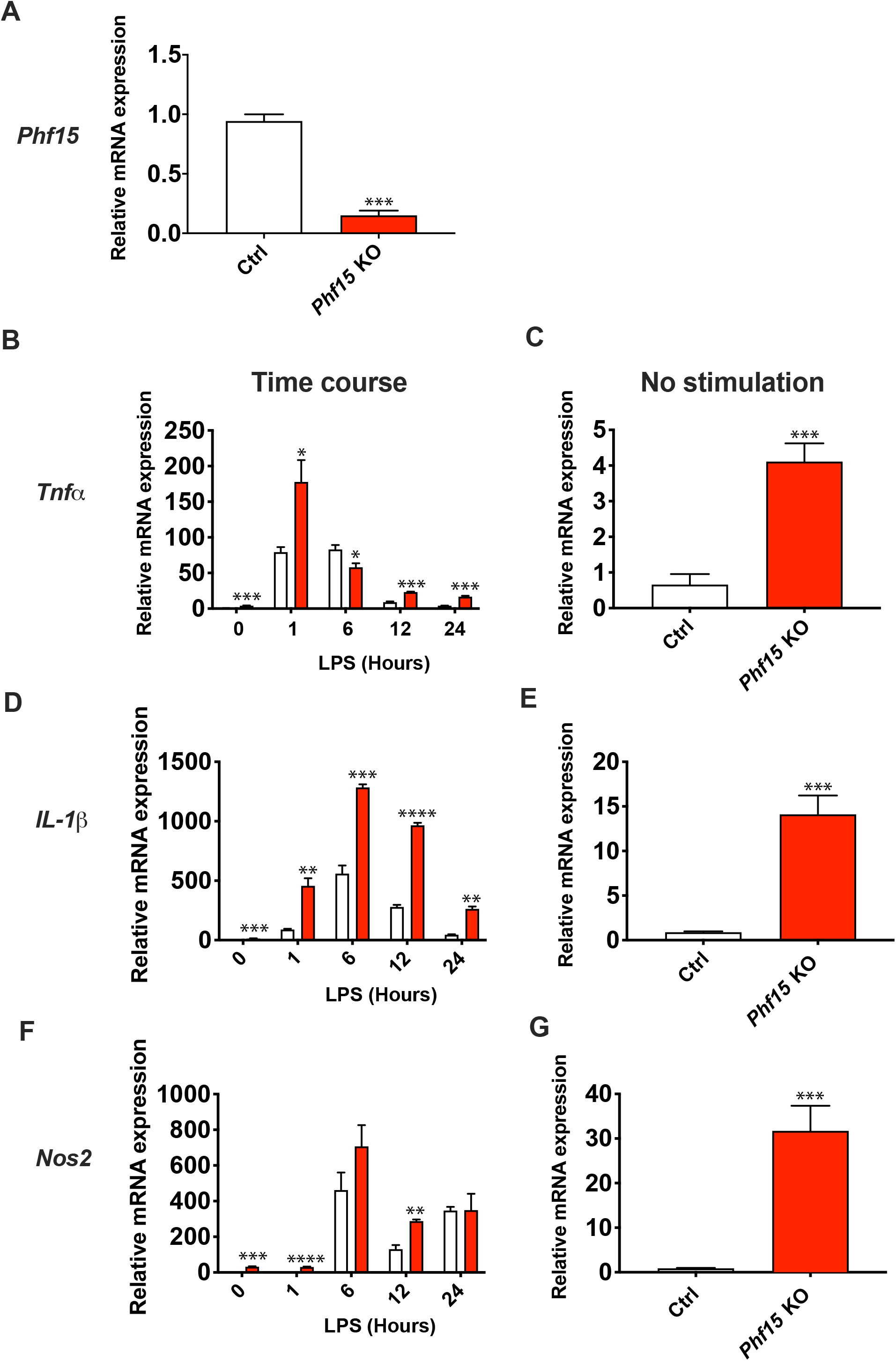
Knockout of *Phf15* increases the magnitude and duration of inflammatory gene expression. (A) Percent reduction in *Phf15* transcript expression in *Phf15* knockout SIM-A9 microglia (*Phf15* KO, red bar) compared to control (Ctrl, open bar). 24-hour time course experiments showing relative mRNA expression levels of *Tnfα* (B), *Il-1β* (D), and *Nos2* (F) after LPS stimulation. No stimulation (0 hr time point or baseline) expression of *Tnfα* (C), *Il-1β* (E) and *Nos2* (G) are also shown. All data are mean ±SEM (n = 3 per condition). Unpaired t-tests between *Phf15* KO and control cells for percent reduction and for individual timepoints: asterisks indicate \**p*<0.05, \*\**p*<0.01, \*\*\**p*<0.001, \*\*\*\**p*<0.0001. LPS, lipopolysaccharide; *Tnfα*, tumor necrosis factor alpha; inducible; *Il-1β*, Interleukin 1 beta; *Nos2*, nitric oxide synthase.

Time course experiments after stimulation of TLR9 with CpG-ODN (Supplementary Figure 4) and TLR3 with Poly(I:C) (Supplementary Figure 5) in *Phf15* KO cells again yielded similar results to LPS stimulation in *Phf15* KO microglial cells, denoting no difference in downstream adapter selectivity and confirming our prior KD results.

Overall, KO of *Phf15* resulted in a more severe phenotype compared to our KD results, increasing the magnitude and prolonging the duration of the microglial inflammatory response. Taken together, our KD and KO results indicate that *Phf15* functions to restrict microglial inflammatory output, regulating the magnitude and duration, as well as, basal inhibition of the inflammatory response.

### Overexpression of *Phf15* in microglia results in a dampened inflammatory response

To further test the role of *Phf15* as a repressor of pro-inflammatory genes, we carried out gain-of-function studies of *Phf15* in SIM-A9 cells. Overexpression (OE) via retroviral delivery of the full-length murine *Phf15* cDNA (Figure 4A) resulted in significantly decreased expression of *Tnfα* at 6 hours (Figure 4B), *1l-1β* (0, 6, 12 hours; Figure 4D) and *Nos2* (0, 6, 24 hours; Figure 4F). Notably, basal levels of both *Il-1β* and *Nos2* were also significantly decreased (Figures 4E, G). Time course experiments following stimulation with CpG-ODN (Supplementary Figure 6) and Poly(I:C) (Supplementary Figure 7) likewise yielded similar results as LPS stimulation of *Phf15* OE microglia, displaying no adapter selectivity.

**Figure 4.**
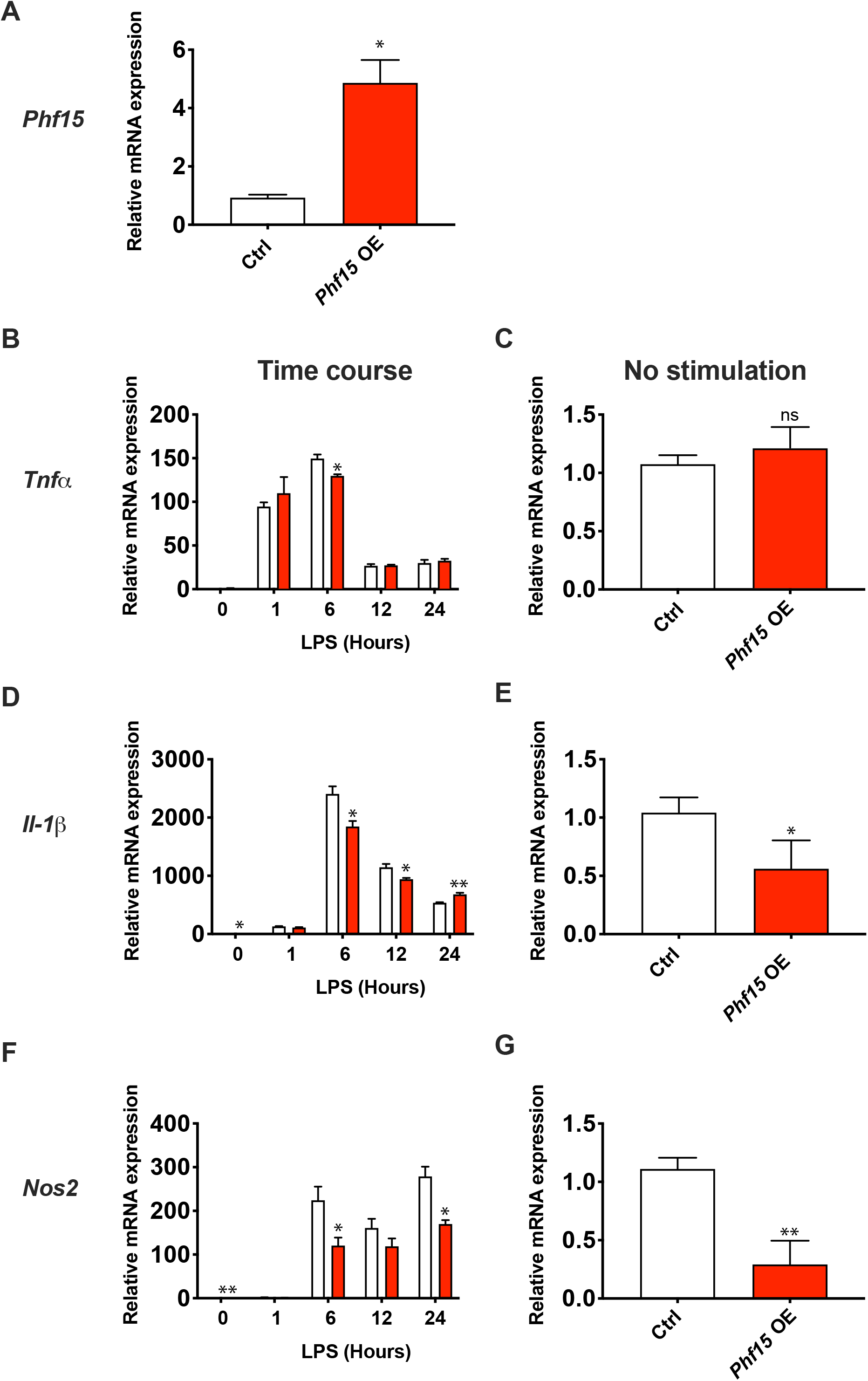
*Phf15* overexpression decreases the microglial inflammatory response. (A) Fold overexpression (OE) of *Phf15* in SIM-A9 microglia (red bar) versus control cells (Ctrl, open bar). 24-hour time course experiments showing relative mRNA expression levels of *Tnfα* (B), *Il-1β* (D), and *Nos2* (F) after LPS stimulation. Baseline (0 hour time point, No stimulation) expression of *Tnfα* (C), *Il-1β* (E), and *Nos2* (G) are displayed separately from time course experiments. All data are mean ±SEM (n = 3 per condition). Unpaired t-tests between *Phf15* OE and control cells for fold-overexpression and for individual time points: asterisks indicate \**p*<0.05, \*\**p*<0.01. LPS, lipopolysaccharide; *Tnfα*, tumor necrosis factor alpha; inducible; *Il-1β*, Interleukin 1 beta; *Nos2*, nitric oxide synthase.

Taken together, our OE results show a dampened microglial inflammatory response, revealing a reciprocal response phenotype compared to our KD and KO experiments. Collectively these results confirm that *Phf15* functions to repress both basal and stimulus-dependent inflammatory gene expressions in microglia.

### Loss of *Phf15* affects global expression of genes involved in antiviral responses and regulation of inflammatory processes

To examine global transcriptional changes as a result of *Phf15* deletion in microglia, we carried out RNA-sequencing (RNA-seq) on *Phf15* KO SIM-A9 cells under no stimulation conditions and 6 hours post LPS stimulation. We chose to examine the no stimulation condition (0 hour time point) based on our KD and KO time course results which showed that baseline is one of the most consistently and strongly deregulated time points. Importantly, elevated or ‘leaky’ pro-inflammatory mediator expression at baseline might result in chronic inflammation leading to neurodegeneration. Similarly, 6 hours after LPS stimulation corresponded to the peak of the transcriptional inflammatory response, with large increases in magnitude for both *Il-1β* and Nos2. Differential gene expression analysis revealed that 466 genes with log2 fold change > 1.5 and *p adj* < 0.01 were up-regulated and 309 genes with log2 fold change < −1.5 and *p adj* < 0.05 were downregulated (Figure 5A). Biological theme enrichment analysis using Metascape^[34]^ on the upregulated genes revealed that the most enriched biological process categories under basal conditions were “response to virus” and “cytokine production” (Figure 5B, C). Under the “response to virus” category, there was significant upregulation of various interferon-stimulated genes (ISGs), for example *Ifit1, Ifit3, Irf7, Isg15, Oas2* and *Oasl2* (Figure 5C). The downregulated genes show more variability in the types of pathways affected, largely involving growth, differentiation and glial cell migration processes (Figure 5A and Supplementary Figure 8A).

**Figure 5.**
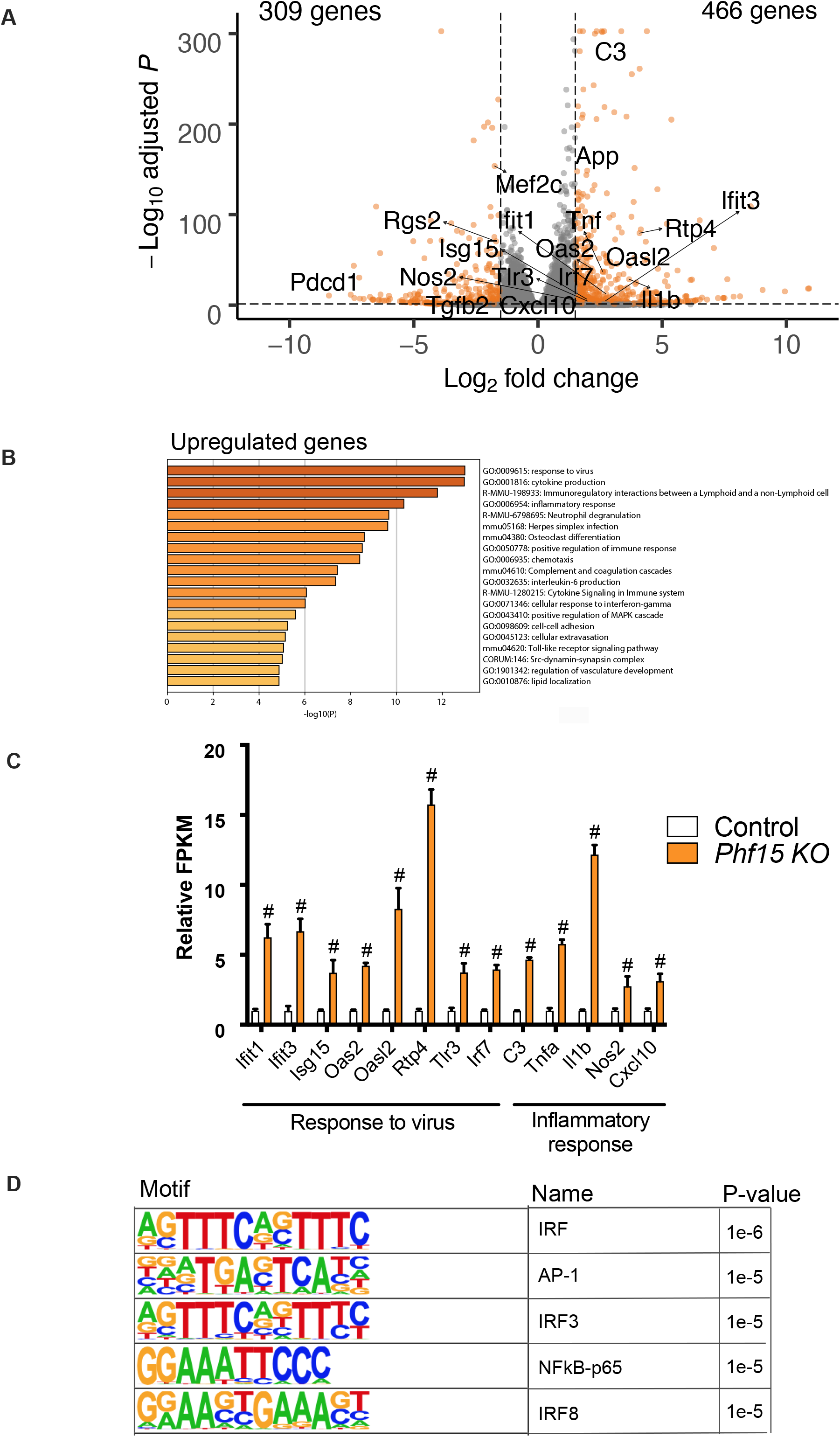
Loss of *Phf15* affects the expression of genes involved in viral response and regulation of inflammatory processes in the absence of immune stimulation. (A) Volcano plot representing the RNA-seq results. Orange dots represent differentially expressed genes in *Phf15* knockout microglia compared to control (upregulated genes at a cutoff of log2fold change > 1.5 and *p adj* < 0.01; downregulated genes at a cutoff of log2fold change < −1.5 and *p adj* < 0.05). (B) GO analysis for significantly upregulated genes showing biological process categories related to response to virus and inflammatory response. (C) Upregulated genes associated with response to virus and inflammatory response in the No stimulation (baseline) condition. Relative FPKM values were obtained by normalizing FPKM values of *Phf15* knockout SIM-A9 microglia to control FPKM values for each gene (n = 3 per condition). Statistics are by DESeq2: asterisks indicate \*\**p*<0.01, #p<0.0001. (D) Top 5 enriched transcription factor binding motifs for the set of upregulated genes in the No stimulation (baseline) condition.

Motif analysis for transcription factor (TF) binding sites enriched in the promoters of the upregulated genes at baseline revealed consensus motifs for Interferon (IFN) stimulated response element (ISRE, IRF binding motif), and motifs for IFN response factor 3 (IRF3) and IRF8 in the top 5 best matches. Also enriched were Activator protein 1 (AP-1) and nuclear factor kappa-light-chain-enhancer of activated B cells p65 subunit (NF-κB-p65) motifs. Both can regulate expression of canonical pro-inflammatory cytokines such as *Tnfα* and Il-1.^[45,46]^ (Figure 5D). Motif enrichment for the set of downregulated genes revealed motifs for Twist-related protein 2 (Twist2) and basic helix–loop–helix (bHLH) MIST1(BHLHA15). Twist2 has been shown to mediate cytokine downregulation after chronic NOD2 (a bacterial peptidoglycan sensor) stimulation^[47]^. MIST1 has been shown to induce and maintain secretory architecture in cells specialized for secretion^[48]^ (Supplementary Figure 8B).

Differential gene expression analysis after 6 hours of LPS stimulation in KO versus control cells revealed 576 up-regulated genes (log2 fold change > 1.5 and *p adj* < 0.01) and 322 down-regulated genes (log2 fold change < −1.5 and *p adj* < 0.05) (Figure 6A). Interestingly, by 6 hours after LPS administration, some of the most enriched biological process categories in KO cells were related to “cytokine secretion” and “immunoregulatory interaction” (Figure 6B and C), denoting a strong increase in magnitude of expression of genes involved in regulating the secretion of pro-inflammatory mediators. The downregulated genes at 6 hours after LPS stimulation in KO cells relative to control again displayed more variability, but did show decreases in biological process categories related to “regulation of defense response” and “cytokine production”, indicating negative regulation of these processes in *Phf15* KO cells compared to control (Supplementary Figure 9A).

**Figure 6.**
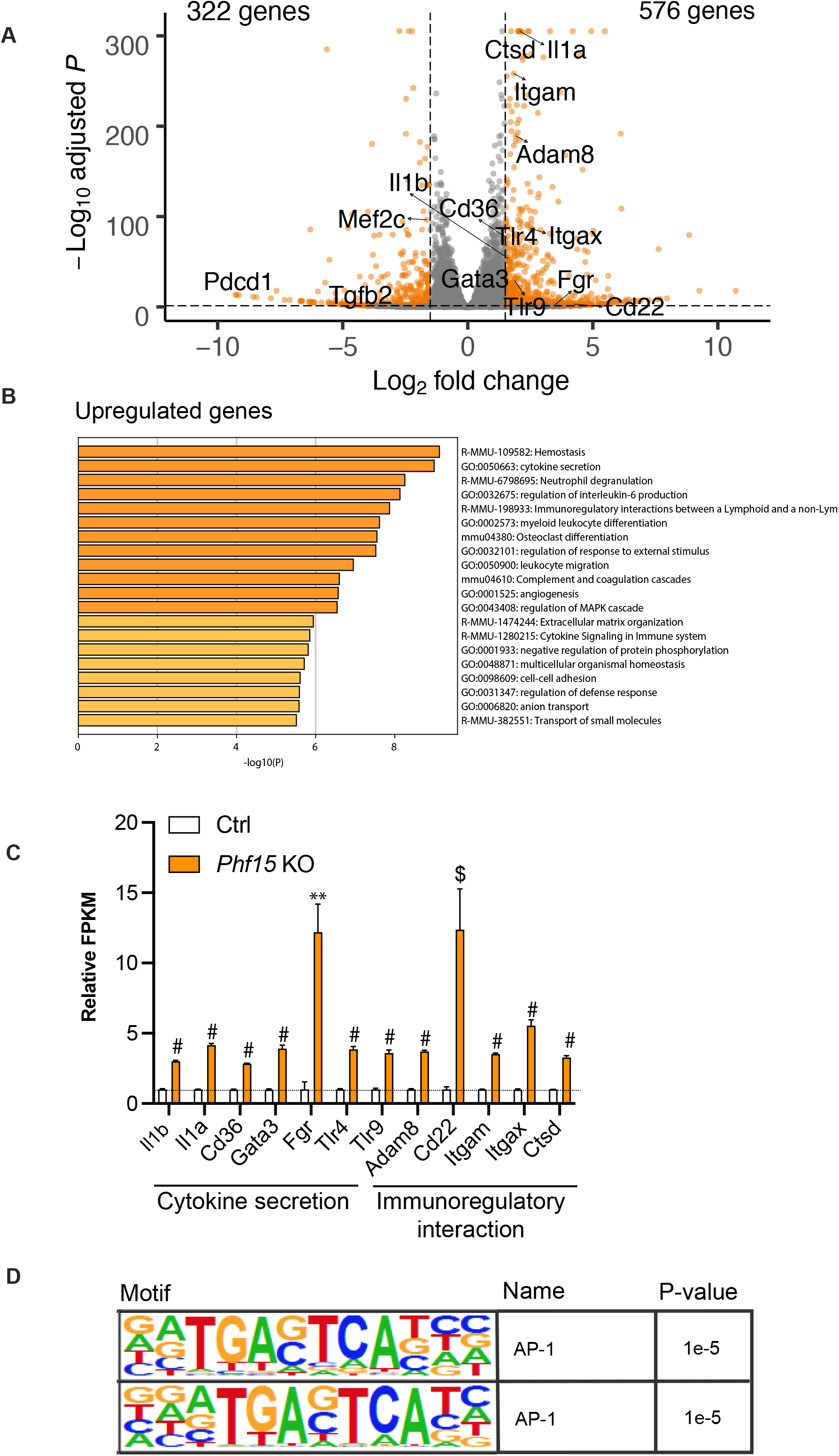
Knockout of *Phf15* affects the expression of genes involved in inflammatory factor secretion and immunoregulatory processes after LPS stimulation. (A) Volcano plot representing the RNA-seq results. Orange dots represent differentially expressed genes in *Phf15* knockout microglia 6 hours after LPS administration compared to control (upregulated genes at a cutoff of log2fold change > 1.5 and *p adj* < 0.01; downregulated genes at a cutoff of log2fold change < −1.5 and *p adj* < 0.05). (B) GO analysis for upregulated genes shows biological process categories associated with cytokine secretion and immunoregulatory interaction. (C) Upregulated genes associated with cytokine secretion and immunoregulatory interaction biological process categories 6 hours post LPS stimulation. Relative FPKM values were obtained by normalizing FPKM values of *Phf15* knockout SIM-A9 microglia to control FPKM values for each gene (n = 3 wells per condition). Statistics are by DESeq2: asterisks indicate \*\**p*<0.01, $p<0.001, #p<0.0001. (D) Transcription factor binding motifs for the set of upregulated genes 6 hours after LPS stimulation are enriched for Activator protein 1 (AP-1).

Motif enrichment analysis for TF binding sites enriched in the promoters of upregulated genes at the 6-hour time point revealed consensus sequences for AP-1, a key regulator of microglia reactivity in inflammation^[49]^(Figure 6D). Motif enrichment for the set of downregulated genes revealed ISRE, such as IRF1 and IRF3 motifs (Supplementary Figure 9B), supporting the observation that there is a negative “regulation of defense response” by 6 hours post stimulation. It is interesting to note that a functional transition from cytokine production to cytokine secretion seems to occur in the 6 hour period after LPS activation.

Taken together, our RNA-seq results confirm that *Phf15* is a repressor of microglial inflammatory gene expression, regulating the antiviral response - specifically, IFN-I-dependent responses - as well as processes related to pro-inflammatory cytokine production and release.

## DISCUSSION

Our results show that *Phf15* inhibits microglial expression of pro-inflammatory mediators under basal and signal-dependent activation, regulating both the magnitude and duration of the inflammatory response. Genetic deletion of *PhfF15* in a microglial cell line followed by stimulation with LPS lead to an exaggerated pro-inflammatory response with increased production of *Tnfα, Il-1β* and *Nos2* over a time course of 24 hours. Importantly, levels of pro-inflammatory factors remained elevated at 24 hours demonstrating a sustained and prolonged response. Consistent with our LPS stimulation of TLR4 results, similar results were obtained after TLR9 and TLR3 activation confirming that *Phf15* is a general negative regulator and controls *both* the MyD88 and TRIF downstream signal transduction pathways (Supplementary Figure 1). Overexpression of *Phf15* showed a dampened microglial inflammatory response, highlighting a reciprocal response phenotype that further supports our loss-of-function results.

Prolonged inflammation can damage surrounding healthy tissue, eventually resulting in neuronal degeneration and loss, and negatively affecting brain function. For example, levels of *Tnfα* are seen to rapidly rise in experimental models of PD and are highly toxic to dopaminergic neurons^[12,13,50]^. Similarly, high levels of TNFα are a hallmark of PD in humans^[51–53]^. Additionally, both TNFα and IL-1β are involved in maintaining proper synaptic plasticity at physiological levels^[54,55]^ and overproduction of these cytokines can result in neuronal death via excitoxicity and cognitive dysfunction^[56,57]^.

Our studies further demonstrate that *Phf15* can regulate both basal and signal-dependent microglial inflammatory gene expression. KD and KO of *Phf15* in microglial cell lines resulted in significantly increased levels of pro-inflammatory cytokine gene expression 1) without stimulation and 2) after immune activation, while OE had the reverse effect. The inflammatory response is a tightly controlled process in immune cells in order to protect against unintended damage to healthy tissue. Even in aged microglia, where production and secretion of pro-inflammatory mediators is generally increased, this process is dependent upon treatment with immune stimulants^[9,58,59]^. Increased pro-inflammatory cytokine gene expression *without* stimulation denotes constitutive or ‘leaky’ expression of inflammatory mediators, simulating a state of low-grade but constant activation. Similarly, hyperresponsiveness to immune stimuli combined with a lack of resolution of the inflammatory response can lead to a state of chronic inflammation. All three can trigger pathological chronic inflammation in the brain which is detrimental to brain function.

Importantly, distinct molecular mechanisms regulate transcriptional control of different phases (‘modules’) of the inflammatory response and it is noteworthy that *Phf15* might be involved in regulating several of these. Basal inflammatory function, for example, is generally regulated by co-repressors such as nuclear receptor co-repressor (NCOR), silencing mediator of retinoid and thyroid receptors (SMRT) and REST co-repressor 1, (RCOR1 or CoREST) that block poised promoters from active transcription, preventing ‘leaky’ expression of primary response genes (e.g., *TNFα*, Type I IFNs, *Il-1β*, etc.)^*[15]*^. Significantly increased inflammatory gene transcription under baseline conditions, as observed in our *Phf15* KD and KO experiments, suggests a loss of this repressive mechanism.

After stimulation by an activating signal, additional mechanisms can maintain quiescence by restraining active transcription. For example, nuclear receptors like peroxisome proliferator-activated receptor-γ (PPARγ), glucocorticoid receptor (GR) and liver X receptors (LXRs) can inhibit the signal-activated exchange of co-repressors for co-activators at poised promoters, inhibiting the initiation of transcription^*[15,17]*^. Lastly, several mechanisms regulate resolution of inflammation at the transcriptional level, including transrepression mechanisms that can remove transcription factors like NF-κB, from inflammatory gene promoters, effectively blocking expression of secondary response genes, that is, genes which require chromatin-modification as well as protein synthesis for their induction (for example *Nos2* and ISGs)^*[8,15,18]*^. Timely resolution of an inflammatory response is crucial in order to limit cellular and tissue damage caused by prolonged or chronic inflammation. Our results suggest that *Phf15* may be involved in regulating all three of the abovementioned mechanisms.

But how might *Phf15* be involved in regulating transcriptional repression of the inflammatory response? PHF15 was first described in embryonic stem cells as an E3 ligase that directly targets Lysine-specific demethylase 1 (LSD1, Kdm1a) - a key demethylase of histone 3 lysine 4 - for degradation^*[22]*^. LSD1 has been identified as a member of the CoREST co-repressor complex^[60,61]^ which is required for transcriptional repression of inflammation in microglia^*[8]*^. We therefore initially hypothesized that increased levels of *Phf15* upon aging might lead to decreased levels of LSD1 and increased microglial inflammatory output. Our results, however, demonstrate that *Phf15 itself* inhibits microglial inflammatory function, thus, its purported mechanism for inhibition is likely not via degradation of LSD1.

Interestingly, the global transcriptional changes caused by *Phf15* deletion are highly similar to age-associated transcriptional changes in microglia that have been previously reported^[9,62,63]^. In particular, a study by Deczkowska et al.^[64]^, found “immune system process” and specifically “response to virus” among the most highly upregulated biological categories for differentially expressed genes in microglia of young (2-month old) versus aged (22-month old) mice, consistent with our results in *Phf15* KO microglia. Notably, a study by Hammond et al.^[63]^, which used single-cell RNAseq to look at microglia profiles throughout the mouse lifespan, found subpopulations in aged (P540) mouse brains which were largely 1) inflammatory, that is, they upregulated *Il-1β, Tnfα* and other cytokines or 2) IFN-I-responsive, upregulating ISGs, particularly Ifit3, Irf7, Isg15, Oasl2, Ifitm3, and Rtp4, compared to younger adult (P100) brains. Similarly, a recent study from the Tabula Muris Consortium which produced a single-cell transcriptomic atlas of 23 tissues and organs across the *Mus musculus* life span, confirmed that microglia in the aged (P540 and P720) brain are enriched for IFN-I-responsive genes and upregulate a similar set of genes including Ifit3, Irf7, Isg15, Oasl2, Ifitm3, and Rtp4^[65]^. The genes upregulated by the interferon-responsive microglia clusters in both these studies are highly similar to those upregulated in our *Phf15* KO cells under basal conditions (see Figure 5A and C). Because ISGs can modulate inflammation^*[24]*^, it is possible that interferon-responsive microglia could play a role in contributing to the inflammatory signature found in the aged brain. Interestingly, among the set of downregulated genes in *Phf15* KO cells at baseline and 6 hours after LPS stimulation, is Myocyte Enhancer Factor 2C (Mef2C). Mef2C is an important checkpoint inhibitor that restrains microglial activation in response to pro-inflammatory insults and is lost in brain aging via IFN-I mediated downregulation^[64,66]^. Thus, an increase in *Phf15* expression in microglia during healthy aging could putatively work to counteract not only microglial activation but increased IFN-I in the aged brain as well.

Notably, a recent study by Readhead et al.,^[67]^ found that several virus species are commonly present in the aged human brain. Among them, human herpesvirus 6A and 7 (HHV-6A and HHV-7) were highly upregulated in the brain of AD patients and were found to modulate host genes associated with AD risk, for example, Amyloid precursor protein (APP) processing. APP is the precursor molecule whose proteolysis forms amyloid-β (Aβ) and formation of Aβ plaques has long been thought of as the driving force behind Alzheimer’s disease^[62]^. Aβ has more recently been found to have antimicrobial properties, conferring increased resistance against infection from both bacteria and viruses^[63]^. APP is among the significantly upregulated genes under basal conditions in our *Phf15* KO cells (log2 fold change =1.492 and *p adj* < 0.0001; see Figure 5A). Upregulation of APP due to loss of *Phf15* in mouse microglia is thus consistent with our data showing *Phf15* regulation of the antiviral microglial response.

Altogether, our results show that *Phf15* is a novel repressor of microglial inflammatory gene expression, regulating both the magnitude and time-to-resolution of the inflammatory response. Importantly, *Phf15* also serves to repress baseline inflammatory output in the absence of immune activation. Putatively, increases in *Phf15* during healthy aging could help counteract brain inflammation and protect brain health.

Future studies will determine the mechanism of action of *Phf15*. For example, the identity of its binding partner proteins, its genome-wide binding sites and associated histone marks to determine the specific gene regulatory regions it interacts with (e.g. active enhancers or promoters). Additionally, studies in *Phf15* KO mice will elucidate whether loss of Phf15-mediated repression of pro-inflammatory factors is sufficient to induce cognitive decline or exacerbate LPS-induced neurotoxicity of dopaminergic neurons in the substantia nigra.

## DECLARATIONS

## Acknowledgements

We thank Prof. Ellen Robey for helpful comments on the manuscript and Wendy Yan for technical assistance.

## Author’s contributions

Designed and performed experiments, analyzed data, and wrote the manuscript: Muroy, SE

Performed experiments and analyzed data: Timblin, GA, Preininger, MK

Performed experiments: Cedillo, P

Designed experiments and wrote the manuscript: Saijo, K.

## Availability of data and materials

Sequencing data will be deposited in NCBI Gene Expression Omnibus (GEO).

## Ethical approval and consent to participate

All procedures were approved by the Animal Care and Use Committee of the University of California, Berkeley (Animal Use Protocol AUP-2017-02-9539).

## Consent for publication

Not applicable.

## Copyright

The authors.

## References

1. Wake H, Moorhouse AJ, Miyamoto A, Nabekura J. Microglia: actively surveying and shaping neuronal circuit structure and function. Trends Neurosci 2013;36:209–17. http://doi.org/10.1016/j.tins.2012.11.007

2. Parkhurst CN, Gan W-B. Microglia dynamics and function in the CNS. Current Opinion in Neurobiology 2010;20:595–600. http://doi.org/10.1016/j.conb.2010.07.002

3. Kettenmann H, Hanisch U-K, Noda M, Verkhratsky A. Physiology of microglia. Physiol Rev 2011;91:461–553. http://doi.org/10.1152/physrev.00011.2010

4. Ransohoff RM, Perry VH. Microglial physiology: unique stimuli, specialized responses. Annu Rev Immunol 2009;27:119–45. http://doi.org/10.1146/annurev.immunol.021908.132528

5. Saijo K, Glass CK. Microglial cell origin and phenotypes in health and disease. Nature Publishing Group 2011;11:775–87. http://doi.org/10.1038/nri3086

6. Nimmerjahn A, Kirchhoff F, Helmchen F. Resting microglial cells are highly dynamic surveillants of brain parenchyma in vivo. Science 2005;308:1314–8. http://doi.org/10.2174/1381612053381620

7. Hanisch U-K, Kettenmann H. Microglia: active sensor and versatile effector cells in the normal and pathologic brain. Nature Neuroscience 2007;10:1387–94. http://doi.org/10.1038/nn1997

8. Saijo K, Winner B, Carson CT, Collier JG, Boyer L, Rosenfeld MG, et al. A Nurr1/CoREST pathway in microglia and astrocytes protects dopaminergic neurons from inflammation-induced death. Cell 2009;137:47–59. http://doi.org/10.1016/j.cell.2009.01.038

9. Mosher KI, Wyss-Coray T. Microglial dysfunction in brain aging and Alzheimer’s disease. Biochem Pharmacol 2014;88:594–604. http://doi.org/10.1016/j.bcp.2014.01.008

10. Salter MW, Stevens B. Microglia emerge as central players in brain disease. Nature Publishing Group 2017;23:1018–27. http://doi.org/10.1038/nm.4397

11. Hickman S, Izzy S, Sen P, Morsett L, Khoury El J. Microglia in neurodegeneration. Nature Publishing Group 2018;21:1359–69. http://doi.org/10.1038/s41593-018-0242-x

12. McCoy MK. Blocking Soluble Tumor Necrosis Factor Signaling with Dominant-Negative Tumor Necrosis Factor Inhibitor Attenuates Loss of Dopaminergic Neurons in Models of Parkinson’s Disease. J Neurosci 2006;26:9365–75. http://doi.org/10.1523/JNEUROSCI.1504-06.2006

13. Sriram K, Matheson JM, Benkovic SA, Miller DB, Luster MI, O’Callaghan JP. Mice deficient in TNF receptors are protected against dopaminergic neurotoxicity: implications for Parkinson’s disease. The FASEB Journal 2002;16:1474–6. http://doi.org/10.1096/fj.02-0216fje

14. Glass CK, Saijo K, Winner B, Marchetto MC, Gage FH. Mechanisms Underlying Inflammation in Neurodegeneration. Cell 2010;140:918–34. http://doi.org/10.1016/j.cell.2010.02.016

15. Medzhitov R, Horng T. Transcriptional control of the inflammatory response. Nature Reviews Immunology 2009;9:692–703. http://doi.org/10.1038/nri2634

16. Smale ST, Natoli G. Transcriptional Control of Inflammatory Responses. Cold Spring Harbor Perspectives in Biology 2014;6:a016261–1. http://doi.org/10.1101/cshperspect.a016261

17. Glass CK, Ogawa S. Combinatorial roles of nuclear receptors in inflammation and immunity. Nature Reviews Immunology 2005;6:44–55. http://doi.org/10.1038/nri1748

18. Glass CK, Saijo K. Nuclear receptor transrepression pathways that regulate inflammation in macrophages and T cells. Nature Publishing Group 2010;10:365–76. http://doi.org/10.1038/nri2748

19. Smale ST, Tarakhovsky A, Natoli G. Chromatin Contributions to the Regulation of Innate Immunity. Annu Rev Immunol 2014;32:489–511. http://doi.org/10.1146/annurev-immunol-031210-101303

20. Bernstein BE, Meissner A, Lander ES. The Mammalian Epigenome. Cell 2007;128:669–81. http://doi.org/10.1016/j.cell.2007.01.033

21. Soreq L, Rose J, Soreq E, Hardy J, Trabzuni D, Cookson MR, et al. Major Shifts in Glial Regional Identity Are a Transcriptional Hallmark of Human Brain Aging. Cell Rep 2017;18:557–70. http://doi.org/10.1016/j.celrep.2016.12.011

22. Han X, Gui B, Xiong C, Zhao L, Liang J, Sun L, et al. Destabilizing LSD1 by Jade-2 promotes neurogenesis: an antibraking system in neural development. Mol Cell 2014;55:482–94. http://doi.org/10.1016/j.molcel.2014.06.006

23. Panchenko MV. Structure, function and regulation of jade family PHD finger 1 (JADE1). Gene 2016;589:1–11. http://doi.org/10.1016/j.gene.2016.05.002

24. Baruch K, Deczkowska A, David E, Castellano JM, Miller O, Kertser A, et al. Aging-induced type I interferon response at the choroid plexus negatively affects brain function. Science 2014;346:89. http://doi.org/10.1126/science.1252945

25. Sparkman NL, Johnson RW. Neuroinflammation Associated with Aging Sensitizes the Brain to the Effects of Infection or Stress. Neuroimmunomodulation 2008;15:323–30. http://doi.org/10.1159/000156474

26. Gabuzda D, Yankner BA. Physiology: Inflammation links ageing to the brain. Nature 2013;497:197–8. http://doi.org/10.1038/nature12100

27. Lynch MA. Age-related neuroinflammatory changes negatively impact on neuronal function. Front Aging Neurosci 2010;1:1–8. http://doi.org/10.3389/neuro.24.006.2009

28. Godbout JP, Chen J, Abraham J, Richwine AF, Berg BM, Kelley KW, et al. Exaggerated neuroinflammation and sickness behavior in aged mice following activation of the peripheral innate immune system. The FASEB Journal 2005;19:1329–31. http://doi.org/10.1096/fj.05-3776fje

29. Stewart SA. Lentivirus-delivered stable gene silencing by RNAi in primary cells. RNA 2003;9:493–501. http://doi.org/10.1261/rna.2192803

30. Sena-Esteves M, Tebbets JC, Steffens S, Crombleholme T, Flake AW. Optimized large-scale production of high titer lentivirus vector pseudotypes. Journal of Virological Methods 2004;122:131–9. http://doi.org/10.1016/j.jviromet.2004.08.017

31. Dobin A, Davis CA, Schlesinger F, Drenkow J, Zaleski C, Jha S, et al. STAR: ultrafast universal RNA-seq aligner. Bioinformatics 2012;29:15–21. http://doi.org/10.1093/bioinformatics/bts635

32. Love MI, Huber W, Anders S. Moderated estimation of fold change and dispersion for RNA-seq data with DESeq2. Genome Biol 2014;15:31–21. http://doi.org/10.1186/s13059-014-0550-8

33. Blighe K, Rana S, Lewis M. EnhancedVolcano: Publication-ready volcano plots with enhanced colouring and labeling [Internet]. https://github.comkevinbligheEnhancedVolcano Available from: https://github.com/kevinblighe/EnhancedVolcano. http://doi.org/10.18129/B9.bioc.EnhancedVolcano

34. Zhou Y, Bin Zhou, Pache L, Chang M, Khodabakhshi AH, Tanaseichuk O, et al. Metascape provides a biologist-oriented resource for the analysis of systems-level datasets. Nat Commun 2019;10:1–10. http://doi.org/10.1038/s41467-019-09234-6

35. Craik FIM, Salthouse TA. The Handbook of Aging and Cognition. 1st ed. Psychology Press; 2011. http://doi.org/10.4324/9780203837665

36. Simen AA, Bordner KA, Martin MP, Moy LA, Barry LC. Cognitive dysfunction with aging and the role of inflammation. Ther Adv Chronic Dis 2011;2:175–95. http://doi.org/10.1177/2040622311399145

37. Zhao J, Bi W, Xiao S, Lan X, Cheng X, Zhang J, et al. Neuroinflammation induced by lipopolysaccharide causes cognitive impairment in mice. Sci Rep 2019;9:1–12. http://doi.org/10.1038/s41598-019-42286-8

38. Glaros TG, Chang S, Gilliam EA, Maitra U, Deng H, Li L. Causes and consequences of low grade endotoxemia and inflammatory diseases. Front Biosci 2013;5:754–65. http://doi.org/10.2741/s405

39. Sandiego CM, Gallezot J-D, Pittman B, Nabulsi N, Lim K, Lin S-F, et al. Imaging robust microglial activation after lipopolysaccharide administration in humans with PET. Proc Natl Acad Sci USA 2015;112:12468–73. http://doi.org/10.1073/pnas.1511003112

40. Chen Y-C, Yip P-K, Huang Y-L, Sun Y, Wen L-L, Chu Y-M, et al. Sequence variants of toll like receptor 4 and late-onset Alzheimer’s disease. PLoS ONE 2012;7:e50771. http://doi.org/10.1371/journal.pone.0050771

41. Brown GC. The endotoxin hypothesis of neurodegeneration. J Neuroinflammation 2019;16:1–10. http://doi.org/10.1186/s12974-019-1564-7

42. Perez-Pardo P, Dodiya HB, Engen PA, Forsyth CB, Huschens AM, Shaikh M, et al. Role of TLR4 in the gut-brain axis in Parkinson’s disease: a translational study from men to mice. Gut 2019;68:829–43. http://doi.org/10.1136/gutjnl-2018-316844

43. Akira S, Takeda K. Toll-like receptor signalling. Nature Reviews Immunology 2004;4:499–511. http://doi.org/10.1038/nri1391

44. Takeda K, Akira S. TLR signaling pathways. Semin Immunol 2004;16:3–9. http://doi.org/10.1016/j.smimm.2003.10.003

45. Chen L-F, Greene WC. Shaping the nuclear action of NF-κB. Nature Reviews Molecular Cell Biology 2004;5:392–401. http://doi.org/10.1038/nrm1368

46. McDonald PP. Transcriptional regulation in neutrophils: teaching old cells new tricks. Adv Immunol 2004;82:1–48. http://doi.org/10.1016/S0065-2776(04)82001-7

47. Zheng S, Hedl M, Abraham C. Twist1 and Twist2 Contribute to Cytokine Downregulation following Chronic NOD2 Stimulation of Human Macrophages through the Coordinated Regulation of Transcriptional Repressors and Activators. J Immunol 2015;195:217–26. http://doi.org/10.4049/jimmunol.1402808

48. Lo H-YG, Jin RU, Sibbel G, Liu D, Karki A, Joens MS, et al. A single transcription factor is sufficient to induce and maintain secretory cell architecture. Genes & Development 2017;31:154–71. http://doi.org/10.1101/gad.285684.116

49. Kaminska B, Mota M, Pizzi M. Signal transduction and epigenetic mechanisms in the control of microglia activation during neuroinflammation. 2016;3:1–13. http://doi.org/10.1016/j.bbadis.2015.10.026

50. Nagatsu T, Sawada M. Inflammatory process in Parkinson’s disease: role for cytokines. Curr Pharm Des 2005;11:999–1016. http://doi.org/10.2174/1381612053381620

51. Qin X-Y, Zhang S-P, Cao C, Loh YP, Cheng Y. Aberrations in Peripheral Inflammatory Cytokine Levels in Parkinson Disease. JAMA Neurol 2016;73:1316–9. http://doi.org/10.1001/jamaneurol.2016.2742

52. Mogi M, Harada M, Riederer P, Narabayashi H, Fujita K, Nagatsu T. Tumor necrosis factor-alpha (TNF-alpha) increases both in the brain and in the cerebrospinal fluid from parkinsonian patients. Neurosci Lett 1994;165:208–10. http://doi.org/10.1016/0304-3940(94)90746-3

53. Boka G, Anglade P, Wallach D, Javoy-Agid F, Agid Y, Hirsch EC. Immunocytochemical analysis of tumor necrosis factor and its receptors in Parkinson’s disease. Neurosci Lett 1994;172:151–4. http://doi.org/10.1016/0304-3940(94)90684-x

54. Rizzo FR, Musella A, De Vito F, Fresegna D, Bullitta S, Vanni V, et al. Tumor Necrosis Factor and Interleukin-1 βModulate Synaptic Plasticity during Neuroinflammation. Neural Plasticity 2018;2018:1–12. http://doi.org/10.1155/2018/8430123

55. Stellwagen D, Malenka RC. Synaptic scaling mediated by glial TNF-α. Nature 2006;440:1054–9. http://doi.org/10.1038/nature04671

56. Takeuchi H, Jin S, Wang J, Zhang G, Kawanokuchi J, Kuno R, et al. Tumor necrosis factor-alpha induces neurotoxicity via glutamate release from hemichannels of activated microglia in an autocrine manner. J Biol Chem 2006;281:21362–8. http://doi.org/10.1074/jbc.M600504200

57. Ye L, Huang Y, Zhao L, Li Y, Sun L, Zhou Y, et al. IL-1β and TNF-α induce neurotoxicity through glutamate production: a potential role for neuronal glutaminase. J Neurochem 2013;125:897–908. http://doi.org/10.1111/jnc.12263

58. Denver P, McClean P. Distinguishing normal brain aging from the development of Alzheimer’s disease: inflammation, insulin signaling and cognition. Neural Regen Res 2018;13:1719–12. http://doi.org/10.4103/1673-5374.238608

59. Njie EG, Boelen E, Stassen FR, Steinbusch HWM, Borchelt DR, Streit WJ. Ex vivo cultures of microglia from young and aged rodent brain reveal age-related changes in microglial function. Neurobiology of Aging 2012;33:195.e1–12. http://doi.org/10.1016/j.neurobiolaging.2010.05.008

60. Foster CT, Dovey OM, Lezina L, Luo JL, Gant TW, Barlev N, et al. Lysine-specific demethylase 1 regulates the embryonic transcriptome and CoREST stability. Mol Cell Biol 2010;30:4851–63. http://doi.org/10.1128/MCB.00521-10

61. Lee MG, Wynder C, Cooch N, Shiekhattar R. An essential role for CoREST in nucleosomal histone 3 lysine 4 demethylation. Nature 2005;437:432–5. http://doi.org/10.1038/nature04021

62. Grabert K, Michoel T, Karavolos MH, Clohisey S, Baillie JK, Stevens MP, et al. Microglial brain region-dependent diversity and selective regional sensitivities to aging. Nature Neuroscience 2016;19:504–16. http://doi.org/10.1038/nn.4222

63. Hammond TR, Dufort C, Dissing-Olesen L, Giera S, Young A, Wysoker A, et al. Single-Cell RNA Sequencing of Microglia throughout the Mouse Lifespan and in the Injured Brain Reveals Complex Cell-State Changes. Immunity 2019;50:253–6. http://doi.org/10.1016/j.immuni.2018.11.004

64. Deczkowska A, Matcovitch-Natan O, Tsitsou-Kampeli A, Ben-Hamo S, Dvir-Szternfeld R, Spinrad A, et al. Mef2C restrains microglial inflammatory response and is lost in brain ageing in an IFN-I-dependent manner. Nat Commun 2017;8:1–13. http://doi.org/10.1038/s41467-017-00769-0

65. The Tabula Muris Consortium, Pisco AO, McGeever A, Schaum N, Karkanias J, Neff NF, et al. A Single Cell Transcriptomic Atlas Characterizes Aging Tissues in the Mouse. 2019;75:645–43. http://doi.org/10.1101/661728

66. Deczkowska A, Amit I, Schwartz M. Microglial immune checkpoint mechanisms. Nature Neuroscience 2018;21:1–8. http://doi.org/10.1038/s41593-018-0145-x

67. Ben Readhead, Haure-Mirande J-V, Funk CC, Richards MA, Shannon P, Haroutunian V, et al. Multiscale Analysis of Independent Alzheimer’s Cohorts Finds Disruption of Molecular, Genetic, and Clinical Networks by Human Herpesvirus. Neuron 2018;:1–27. http://doi.org/10.1016/j.neuron.2018.05.023

