## Supplemental material for "*Phf15*—a novel transcriptional repressor regulating inflammation in mouse microglia"

**Supplementary Figures 1-9 and Supplementary Table 1.**

**Supplementary Figure 1. Schematic of TLR4, TLR9 and TLR3 signal transduction pathways.** TLR4, activated by LPS, uses *both* the MyD88 and TRIF downstream adapters to transduce its signaling cascade, leading to transcription of canonical pro-inflammatory factors like TNFα, IL-1β and Nos2 as well as, the type I interferons (IFNs). TLR9, activated by CpG ODN, uses the MyD88 adapter, while TLR3, stimulated here by Poly(I:C), uses TRIF.
TLR4, Toll-like receptor 4; LPS, lipopolysaccharide; MyD88, Myeloid differentiation primary response 88; TRIF, TIR-domain-containing adapter-inducing interferon-β; TLR9, Toll-like receptor 9; CpG ODN, CpG Oligodeoxynucleotide; TLR3, Toll-like receptor 3; Poly(I:C), Polyinosinic:polycytidylic acid; AP-1, Activator protein 1; NF-kB, nuclear factor kappa-light-chain-enhancer of activated B cells; TNFα, tumor necrosis factor alpha; IL-1β, Interleukin 1 beta*;* IL-6, Interleukin 6; IRF3, Interferon regulatory factor 3; IRF7, Interferon regulatory factor 7; IFNs, interferons.

**Supplementary Figure 2. Knockdown of *Phf15* increases the magnitude of the microglial inflammatory response after TLR9 stimulation.** Time course experiments showing relative mRNA expression levels of *Tnfα* (A) and *Nos2* (C) after CpG ODN stimulation of *Phf15* knockdown microglial SIM-A9 cells compared to shCtrl (control scrambled shRNA). *Tnfα* and *Nos2* expression at time point 0 from the time course experiments are displayed separately in (B) and (D), respectively. Data are mean ± SEM (n = 3 per condition). Unpaired t-tests for *shPhf15-1* or *shPhf15-2* and shCtrl cells for individual timepoints: asterisks indicate *$p$<0.05, **$p$<0.01, ***$p$<0.001,****$p$<0.0001. Knockdown efficiency for cell lines shPhf15-1 and shPhf15-2 compared to shCtrl is shown in Figure 2a.
CpG ODN, CpG Oligodeoxynucleotide; *Tnfα*, tumor necrosis factor alpha; *Nos2*, nitric oxide synthase, inducible.

**Supplementary Figure 3. Knockdown of *Phf15* increases the magnitude of the microglial inflammatory response after TLR3 activation.** 24-hour time course experiments showing relative mRNAexpression levels of *Tnfα* (A) and *Nos2* (C) after Poly(I:C) stimulation of *Phf15* knockdown microglial SIM-A9 cells compared to shCtrl (control scrambled shRNA). *Tnfα* and *Nos2* expression at time point 0 from the time course experiments are displayed separately in (B) and (D), respectively. Data are mean ± SEM (n = 3 per condition). Unpaired t-tests for *shPhf15-1* or *shPhf15-2* and shCtrl cells for individual timepoints: asterisks indicate *$p$<0.05, **$p$<0.01, ***$p$<0.001,****$p$<0.0001.
Knockdown efficiency for cell lines sh*Phf15*-1 and sh*Phf15*-2 compared to shCtrl is shown in Figure 2a.
Poly(I:C), Polyinosinic:polycytidylic acid; *Tnfα*, tumor necrosis factor alpha; *Nos2*, nitric oxide synthase, inducible.

**Supplementary Figure 4. Knockout of *Phf15* followed by TLR9 stimulation increases the magnitude and duration of inflammatory gene expression.** 24-hour time course experiments showing relative mRNA expression levels of pro-inflammatory factors, *Tnfα (A), Il-1β (C),* and *Nos2* (E) after CpG ODN stimulation. *Tnfα, Il-1β* and *Nos2* expression at time point 0 from the time course experiments are displayed separately in (B), (D) and (F), respectively. All data are mean ± SEM (n = 3 per condition). Unpaired t-tests between *Phf15* KO and control cells for percent reduction and for individual timepoints: asterisks indicate *$p$<0.05, **$p$<0.01, ***$p$<0.001,****$p$<0.0001.

Percent reduction in *Phf15* transcript expression in *Phf15* knockout SIM-A9 microglia compared to control is shown in Figure 3a.

CpG ODN, CpG Oligodeoxynucleotide; *Tnfα*, tumor necrosis factor alpha; inducible; *Il-1β*, Interleukin 1 beta; *Nos2*, nitric oxide synthase.

**Supplementary Figure 5. Knockout of *Phf15* followed by TLR3 activation increases expression of *Il-1β*.** (A) 24-hour time course experiment showing relative mRNA expression levels of *Il-1β* after Poly(I:C) stimulation. *Il-1β* expression at time point 0 from the time course experiments is displayed separately in (B). All data are mean ± SEM (n = 3 per condition). Unpaired t-tests between *Phf15* knockout and control cells for individual timepoints: asterisks indicate *$p$<0.05, **$p$<0.01, ****$p$<0.0001.

Percent reduction in *Phf15* transcript expression in *Phf15* knockout SIM-A9 microglia compared to control is shown in Figure 3a.

CpG ODN, CpG Oligodeoxynucleotide; *Il-1β*, Interleukin 1 beta.

**Supplementary Figure 6. *Phf15* overexpression dampens the microglial inflammatory response after TLR9 stimulation.**

24-hour time course experiments showing relative mRNA expression levels of *Tnfα* (A), *Il-1β* (C), and *Nos2* (E) after CpG ODN stimulation. *Tnfα, Il-1β* and *Nos2* expression at time point 0 from the time course experiments are displayed separately in (B), (D) and (F), respectively. All data are mean ± SEM (n = 3 per condition). Unpaired t-tests between *Phf15* OE and control cells for fold-overexpression and for individual timepoints: asterisks indicate *$p$<0.05, ****$p$<0.0001.

Fold overexpression (OE) of *Phf15* in SIM-A9 microglia compared to control cells is shown in Figure 4a.

CpG ODN, CpG Oligodeoxynucleotide; *Tnfα*, tumor necrosis factor alpha; *Il-1β*, Interleukin 1 beta; *Nos2*, nitric oxide synthase, inducible.

**Supplementary Figure 7. *Phf15* overexpression dampens the microglial inflammatory response after TLR3 activation.**

24-hour time course experiments showing relative mRNA expression levels *Tnfα* (A), *Il-1β (C)*, and *Nos2* (E) after Poly(I:C) stimulation. *Tnfα, Il-1β* and *Nos2* expression at time point 0 from the time course experiments are displayed separately in (B), (D) and (F), respectively. All data are mean ± SEM (n = 3 per condition). Unpaired t-tests between *Phf15* OE and control cells for fold-overexpression and for individual timepoints: asterisks indicate *$p$<0.05, **$p$<0.01.

Fold overexpression of *Phf15* in SIM-A9 microglia compared to control cells is shown in Figure 4a.

Poly(I:C), Polyinosinic:polycytidylic acid; *Tnfα*, tumor necrosis factor alpha; *Il-1β*, Interleukin 1 beta; *Nos2*, nitric oxide synthase, inducible.

**Supplementary Figure 8. Loss of *Phf15* leads to downregulation of genes involved in growth, differentiation and glial cell migration processes under baseline conditions.** RNA-seq was performed on *Phf15* knockout SIM-A9 microglia under No stimulation conditions shown in Figure 5. (A) GO analysis

for biological process categories for the set of significantly downregulated genes in *Phf15* knockout SIM-A9 microglia is shown. Biological categories center around various cell growth and differentiation processes and glial cell migration. (B) Enriched transcription factor binding motifs for the set of downregulated genes in the No stimulation (baseline) condition include Twist2 and MIST1/BHLHA15.
Twist2, Twist-related protein 2; MIST1/BHLHA15, basic helix–loop–helix 15.

**Supplementary Figure 9. Knockout of *Phf15* leads to downregulation of genes involved in regulation of defense response and cytokine production 6 hours after LPS stimulation.** RNA-seq was performed on *Phf15* knockout SIM-A9 microglia after LPS stimulation shown in Figure 6. (A) GO analysis for biological process categories for the set of significantly downregulated genes in *Phf15* knockout SIM-A9 microglia is shown. (B) Enriched transcription factor binding motifs for the set of downregulated genes after LPS stimulation include ISRE, such as IRF1 and IRF3 binding motif.
ISRE, Interferon (IFN) stimulated response element; IRF1, IFN response factor 1; IRF3, IFN response factor 3.

**Supplementary Table 1. List of qPCR primers**

| Gene | Direction | Sequence |
|---|---|---|
| *Phf15* | + | TCAGCATCAAGATGTTCCAAACT |
| | - | TGAGCTGGTATGAATCTGGGA |
| *Tnfα* | + | CCCTCACACTCAGATCATCTTCT |
| | - | GCTACGACGTGGGCTACAG |
| *Nos2* | + | GTTCTCAGCCCAACAATACAAGA |
| | - | GTGGACGGGTCGATGTCAC |
| *Il-1β* | + | CGTGGACCTTCCAGGATGAG |
| | - | CGTCACACACCAGCAGGTTA |
| *Hprt* | + | TCAGTCAACGGGGGACATAAA |
| | - | GGGGCTGTACTGCTTAACCAG |

*Phf15*, PHD finger protein 15; *Tnfα*, tumor necrosis factor alpha; *Nos2*, nitric oxide synthase, inducible; *Il-1β*, Interleukin 1 beta; *Hprt*, Hypoxanthine Phosphoribosyltransferase.

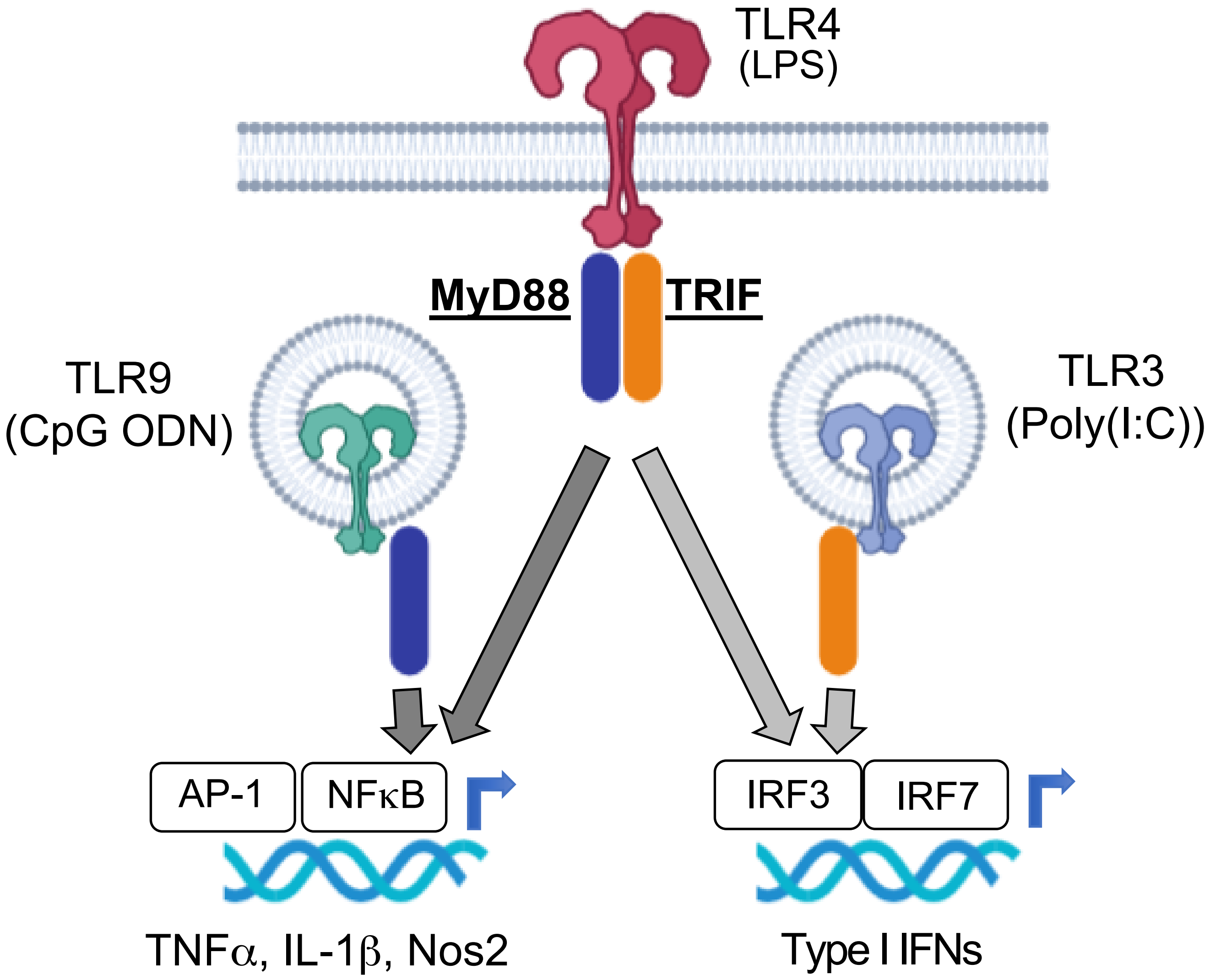
TLR4
(LPS)
MyD88
TRIF
TLR9
(CpG ODN)
TLR3
(Poly(I:C))
AP-1
NFκB
IRF3
IRF7
TNFα, IL-1β, Nos2
Type I IFNs

Sup. Fig. 1

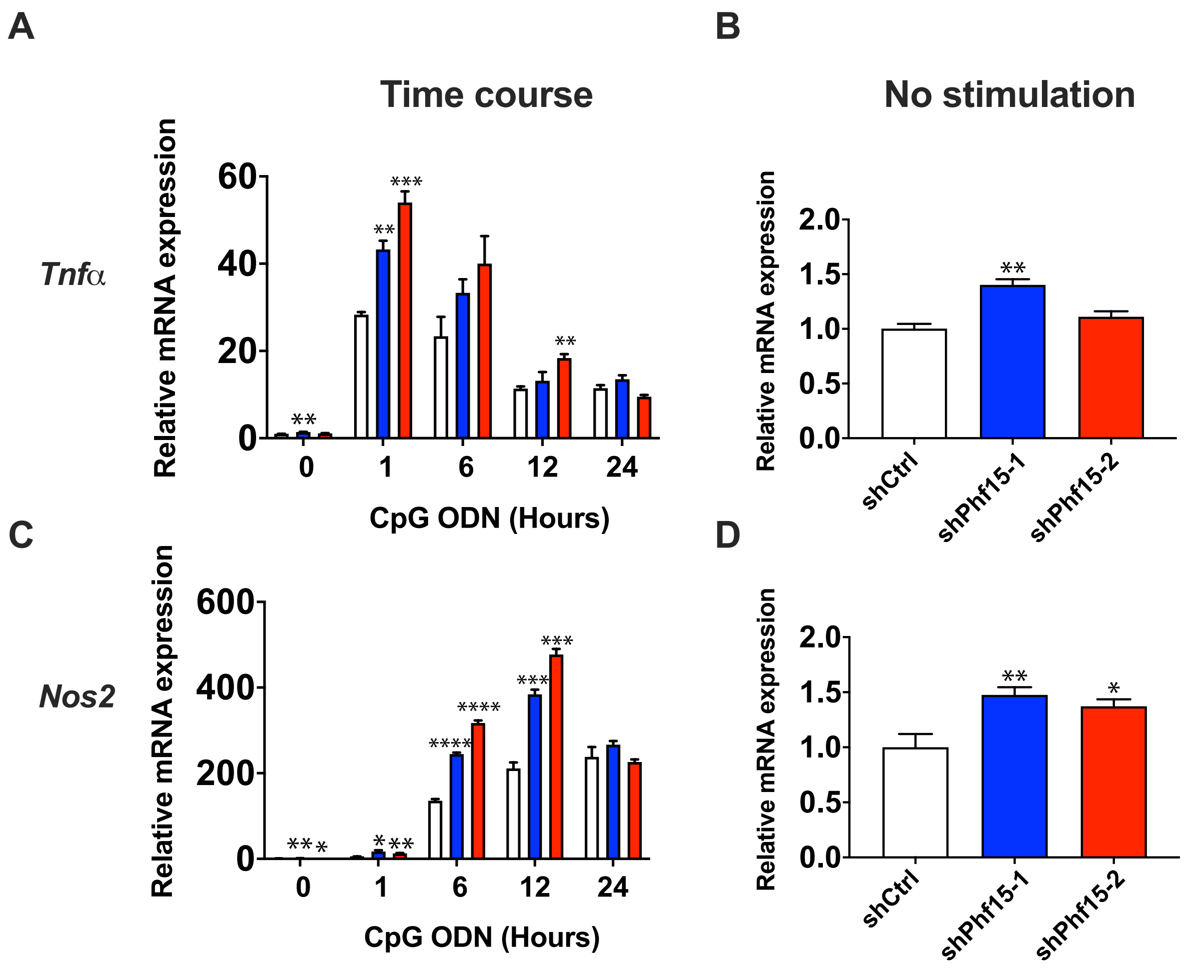
A
B
Time course
No stimulation
Tnfα
Nos2
Relative mRNA expression
CpG ODN (Hours)
shCtrl
shPhf15-1
shPhf15-2
C
D

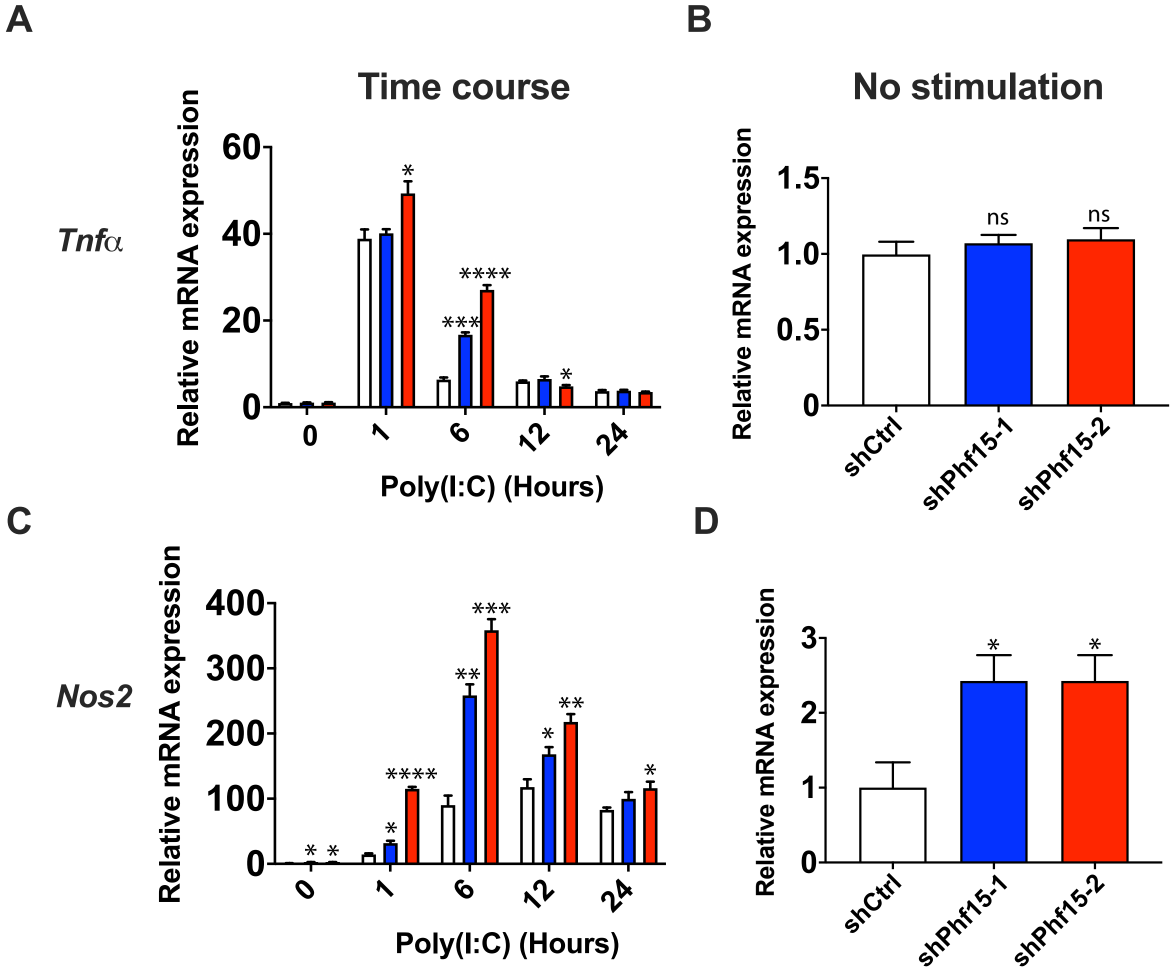
A
B
Time course
No stimulation
Tnfα
Relative mRNA expression
60
40
20
0
*
***
****
*
0
1
6
12
24
Poly(I:C) (Hours)
1.5
1.0
0.5
0
ns
ns
shCtrl
shPhf15-1
shPhf15-2
C
D
Nos2
400
300
200
100
0
****
**
***
**
*
3
2
1
Relative mRNA expression

Sup. Fig. 3

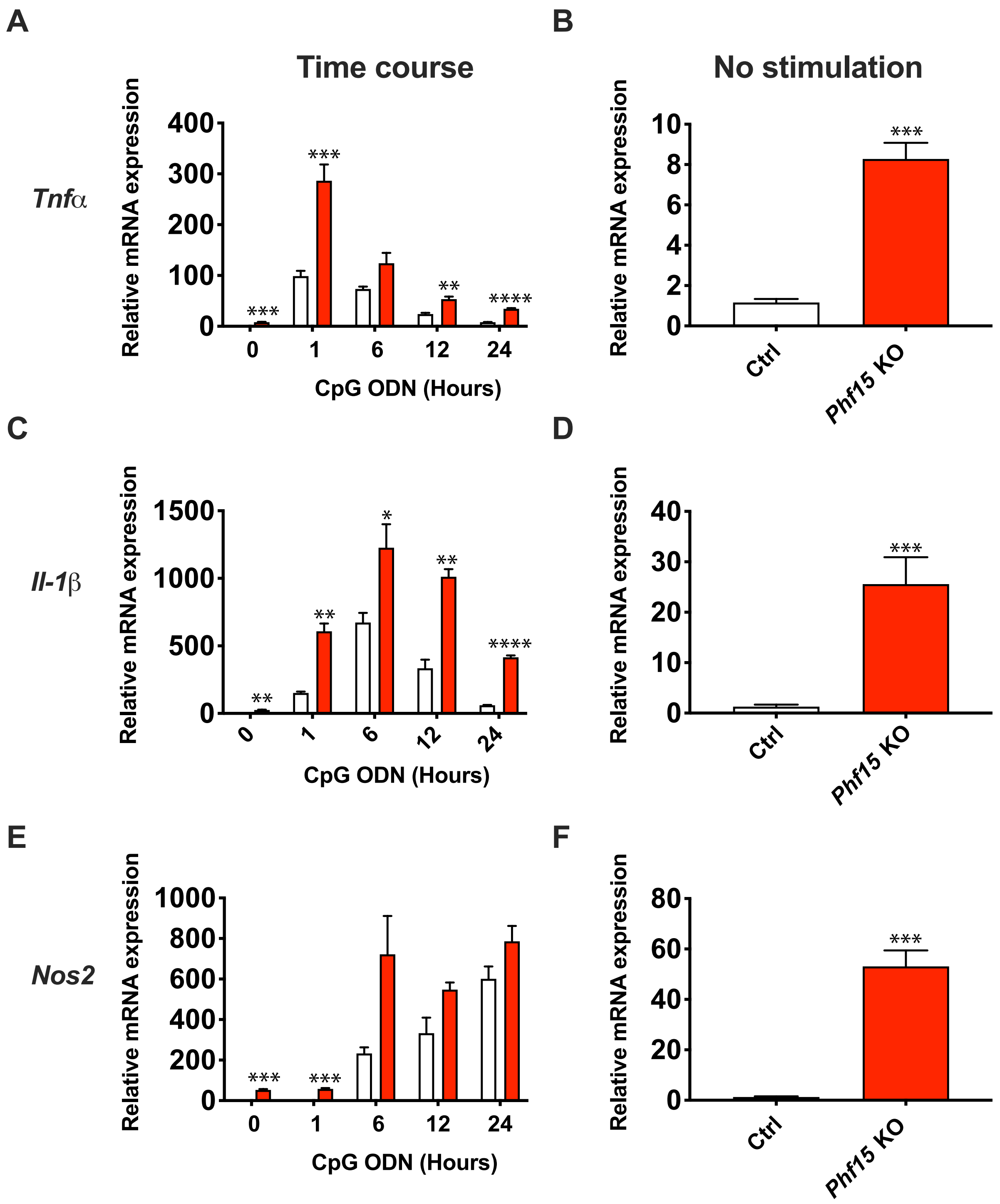
A
B
Time course
No stimulation
Tnfα
Relative mRNA expression
400
300
200
100
0
***
***
**
****
0
1
6
12
24
CpG ODN (Hours)
10
8
6
4
2
0
***
Ctrl
Phf15 KO
C
D
Il-1β
1500
1000
500
0
**
**
*
**
****
40
30
20
10
E
F
Nos2
1000
800
600
400
200
***
80
60
40
20

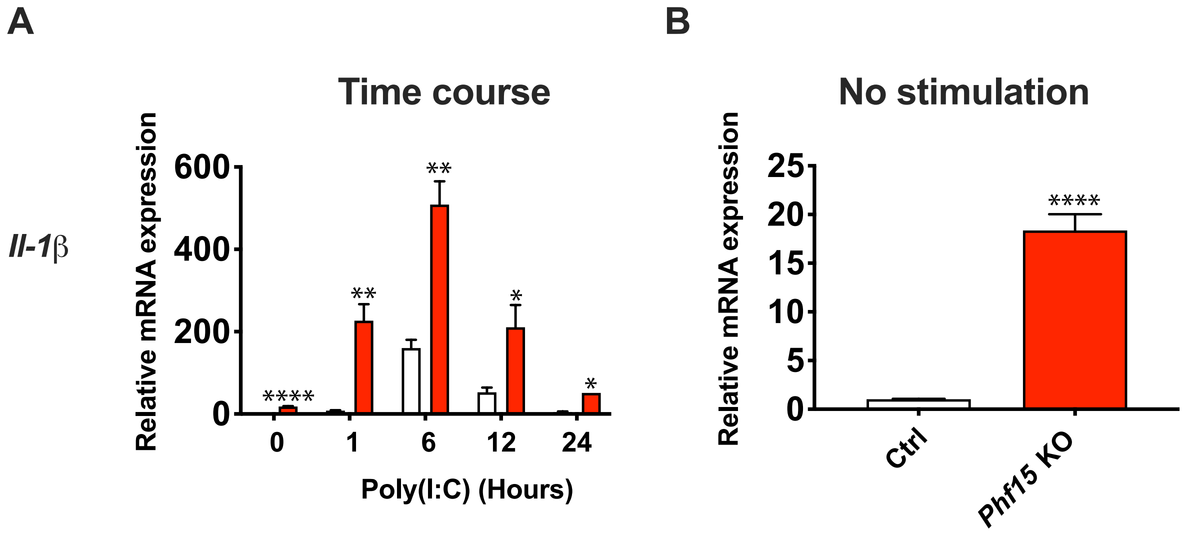
A
B
Il-1β
Time course
Relative mRNA expression
600
400
200
0
****
**
**
*
*
0
1
6
12
24
Poly(I:C) (Hours)
No stimulation
Relative mRNA expression
25
20
15
10
5
0
****
Ctrl
Phf15 KO

Sup. Fig. 5

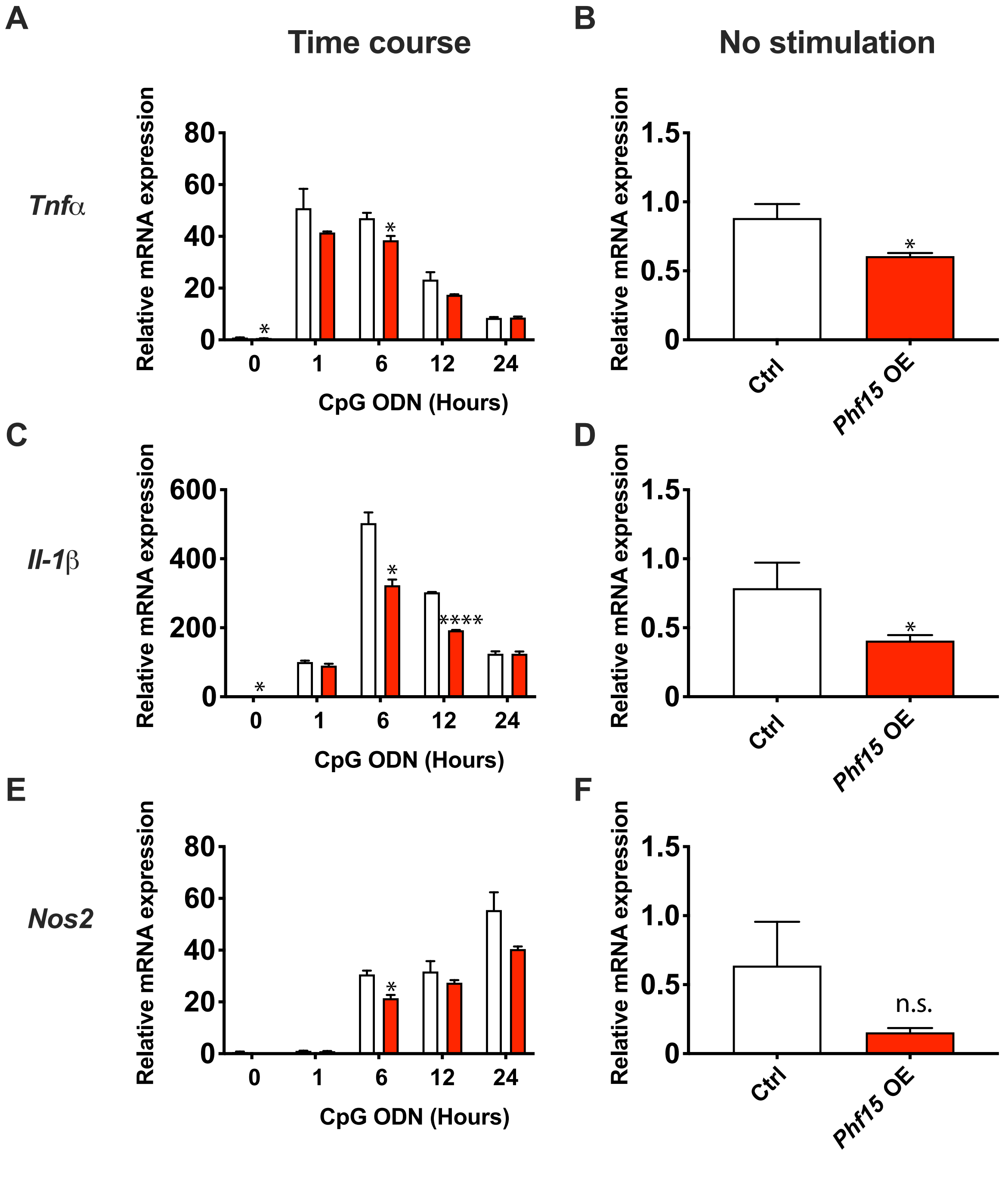

A
Time course
B
No stimulation
Tnfα
Relative mRNA expression
80
60
40
20
0
*
*
0
1
6
12
24
CpG ODN (Hours)
Relative mRNA expression
1.5
1.0
0.5
0
*
Ctrl
Phf15 OE
C
D
Il-1β
Relative mRNA expression
600
400
200
0
*
*
****
0
1
6
12
24
CpG ODN (Hours)
Relative mRNA expression
1.5
1.0
0.5
0
*
Ctrl
Phf15 OE
E
F
Nos2
Relative mRNA expression
80
60
40
20
0
*
0
1
6
12
24
CpG ODN (Hours)
Relative mRNA expression
1.5
1.0
0.5
0
n.s.
Ctrl
Phf15 OE


Sup. Fig. 6

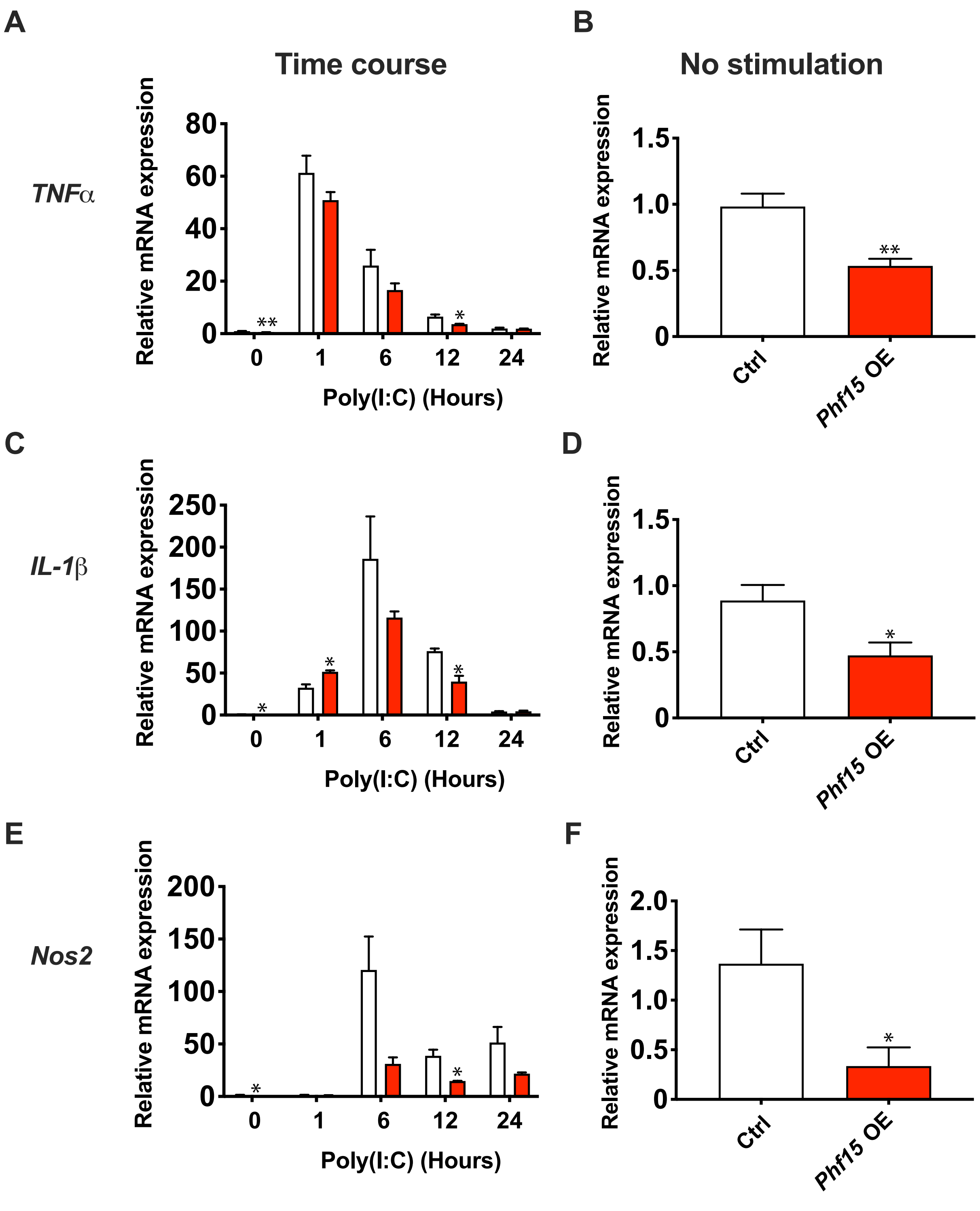

A
Time course
TNFα
Relative mRNA expression
80
60
40
20
0
**
*
0
1
6
12
24
Poly(I:C) (Hours)
B
No stimulation
Relative mRNA expression
1.5
1.0
0.5
0
**
Ctrl
Phf15 OE
C
IL-1β
Relative mRNA expression
250
200
150
100
50
0
*
*
*
0
1
6
12
24
Poly(I:C) (Hours)
D
Relative mRNA expression
1.5
1.0
0.5
0
*
Ctrl
Phf15 OE
E
Nos2
Relative mRNA expression
200
150
100
50
0
*
*
0
1
6
12
24
Poly(I:C) (Hours)
F
Relative mRNA expression
2.0
1.5
1.0
0.5
0
*
Ctrl
Phf15 OE

**A**

### Downregulated genes

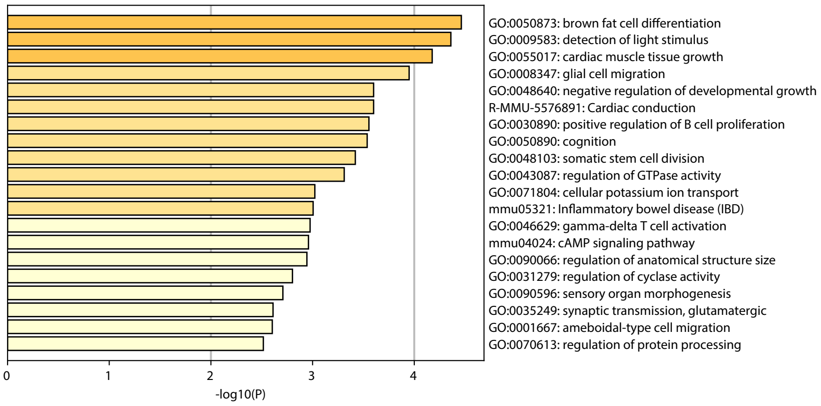


**B**

| Motif | Name | P-value |
|---|---|---|
| 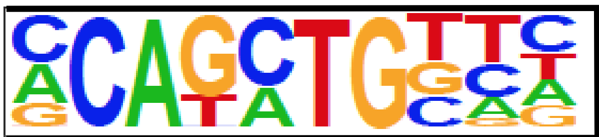 | Twist2 | 1e-4 |
| 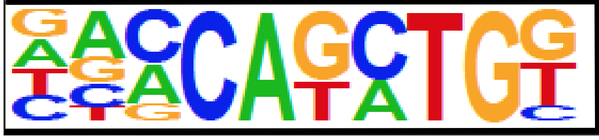 | BHLHA15 | 1e-4 |

**A**

### Downregulated genes

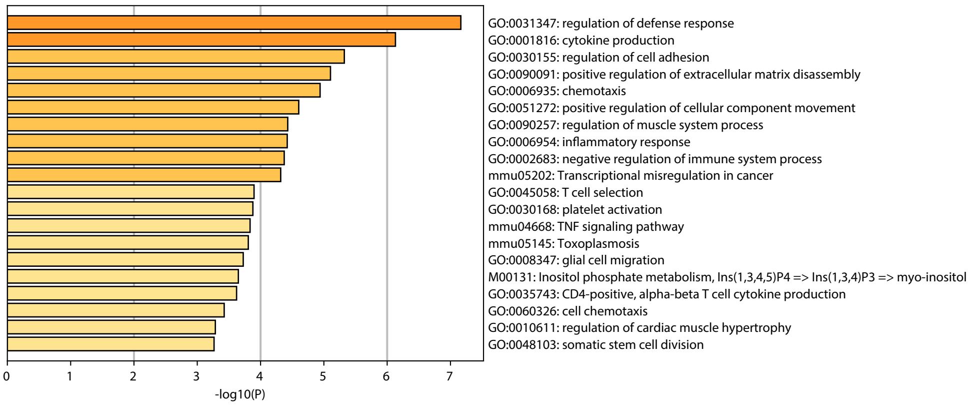


**B**

| Motif | Name | P-value |
|---|---|---|
| 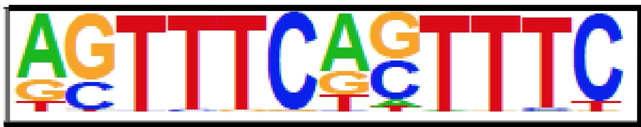 | IRF | 1e-8 |
| 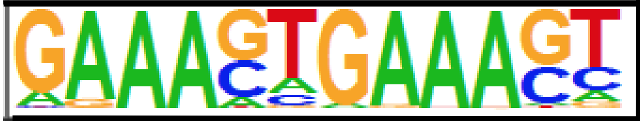 | IRF1 | 1e-5 |
| 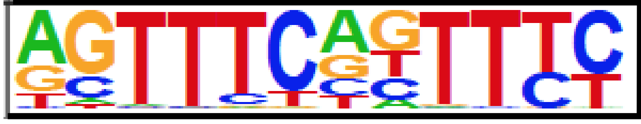 | IRF3 | 1e-5 |
